# Single-cell copy number alteration signature analysis reveals masked patterns and potential biomarkers for cancer

**DOI:** 10.1101/2025.03.02.641098

**Authors:** Chenxu Wu, Ziyu Tao, Wen Chen, Nan Wang, jinyu Wang, jiayu Shen, Tao Wu, Die Qiu, Kaixuan Diao, Xiangyu Zhao, Tianzhu Lu, Lin Zhang, Weiliang Wang, Xinxing Li, Xinxiang Li, Xiaopeng Xiong, Xue-Song Liu

## Abstract

Copy number alteration (CNA) is a major type of cancer genome alteration that drives cancer progression. CNA signature analysis can reveal underlying etiology and provide biomarkers for cancer treatment, and existing CNA signature analyzes are all performed using bulk tissue samples. However CNA usually affect large proportion of genome, and the CNA profile of bulk sample does not reflect the actual CNA profiles of the individual cancer cells of the sample, especially in tumors with high heterogeneity, such as hepatocellular carcinoma (HCC). Furthermore, the evolutionary trajectory of CNA mutational processes still remain elusive. Here we build a method to comprehensively analyze the CNA signatures of HCC from single-cell and bulk sample perspective, revealing patterns and potential noise signals from the usually performed bulk tissue CNA signature analysis. Single-cell signature analysis delineated the evolutionary trajectory of HCC CNA signatures, and different CNA signatures consistently emerge in different HCC evolution stages. Single-cell CNA signatures show robust performance in patient prognosis and drug sensitivity prediction. This work not only reveals specific considerations in analyzing CNA signature derived from bulk tissue but also depicts CNA evolution process and provides potential biomarkers for the prognosis and treatment of HCC patients.

**Highlight:** Single-cell analysis reveals CNA signatures masked in bulk tissue.

Single-cell analysis delineates the evolutionary trajectory of CNA signature.

Small CNAs occur early and large CNAs happens late in HCC evolution.

Single-cell CNA signatures show robust performance in guiding cancer clinical treatment.

## Introduction

Genomic instability is one of the hallmarks of cancer^1^, encompassing changes ranging from single nucleotide alterations to whole chromosome modifications. Numerous studies have demonstrated the roles of single base substitutions (SBS), small insertions and deletions (INDEL), structural variations (including translocations/inversions), and copy number alterations in the process of cancer development ^2–4^. Genomic DNA alteration signatures are recurring genomic patterns that are the imprints of mutagenic processes accumulated over the lifetime of cancer cell^5^. SBS signature analysis has been extensively studied and represents a prototype for other types of signature study^5^. While analyzing copy number signature is more intricate than SBS signatures. Pan-cancer CNA signature studies have been carried out^6–9^, and these studies indicate that genomic alteration signatures can provide information on mutational processes, and also serve as biomarkers for precision medicine in cancer^10–15^.

Current research on CNA signature is predominantly focused on bulk tissue samples. In the detection of SBS, bulk tissue is capable of preserving the SBS mutation status of the majority of subclones. In contrast, CNA usually affect large proportion of genome DNA, and the final CNA profile of bulk tissue is usually not the linear combination of each subclone’s CNA state (Figure 1 as example). For tumors with low heterogeneity, sequencing of bulk tissue effectively captures the variations in copy numbers. However, for tumors with high heterogeneity, like hepatocellular carcinoma (HCC)^16–18^, sequencing of bulk tissue only reflects the averaged CNA across all tumor subclones^19,20^, failing to accurately reveal the true patterns of CNA.

**Figure 1.**
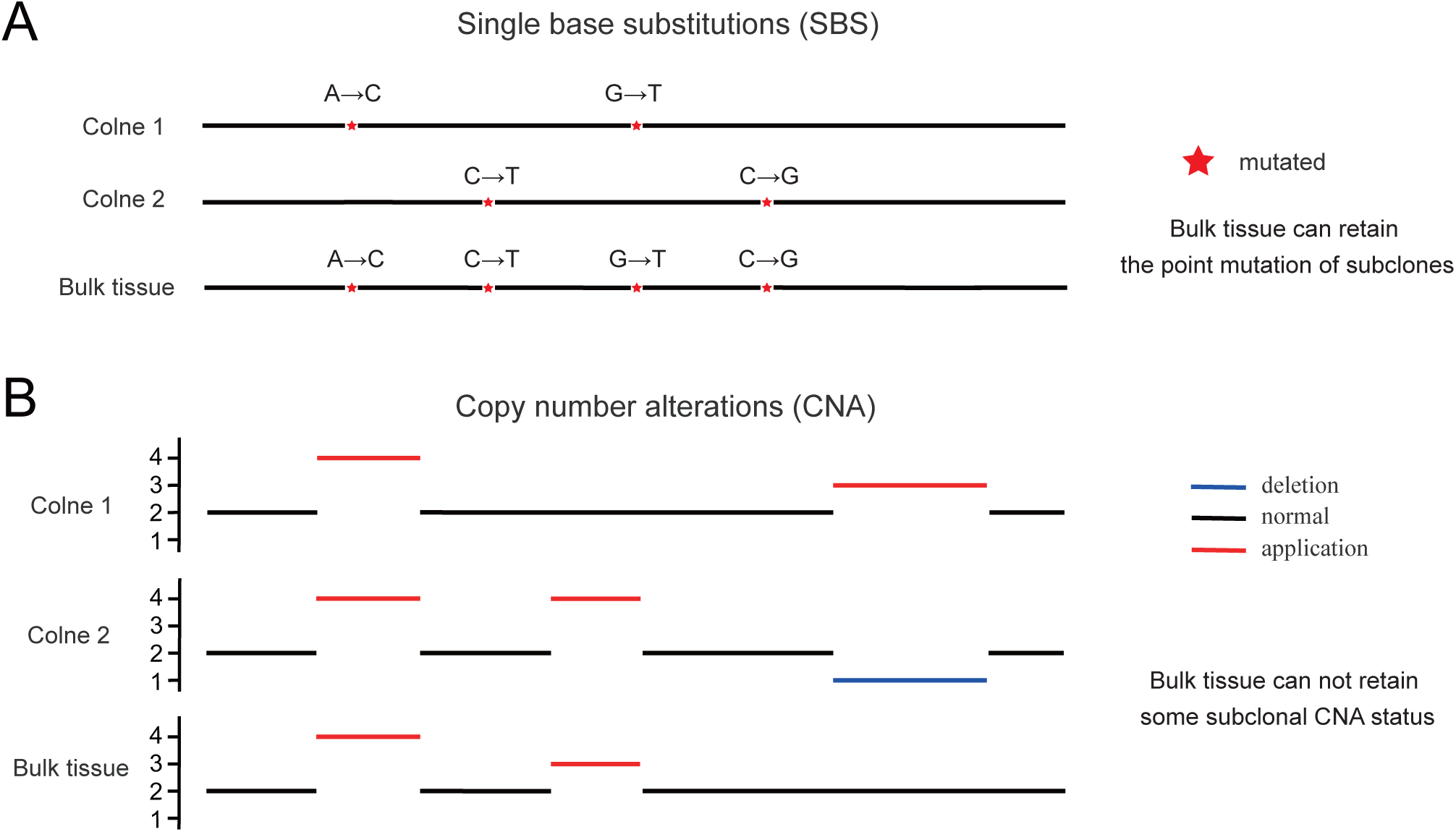
CNA profiles of some subclones may not be reflected in the CNA profiles of bulk tissue. A, Schematic diagram of SBS in different cancer subclones. SBS between different subclones do not affect each other at the bulk tissue level. B, Schematic diagram of CNA in subclones of different cells. The CNA profile of bulk tissue can not retain some subclonal CNA status.

The models of tumor evolution have been a contentious subject in tumor biology^21,22^. Gao et al’s model of punctuated copy number evolution suggests that copy number aberrations are primarily acquired during brief peaks of genomic crisis, followed by stable clonal expansion^23^. Guo et al^24^ introduced a model of dual-phase copy number evolution in HCC, indicating the coexistence of gradual and punctuated evolution. Although these tumor evolution models have revealed the occurrence rate of CNAs during tumor progression, they have not clarified which CNA patterns or signatures appear in the early stages of a tumor and which appear in the later stages. The evolutionary trajectory of CNA signatures still remain elusive.

In this study, we collected scDNA-seq data from 1222 cells of 10 HCC patients and developed a versatile tool for analyzing CNA signatures that apply to both single-cell and bulk tissue samples. This tool enabled us to systematically analyze the CNA signatures in HCC at the single-cell level for the first time. Comparing CNA signatures derived from single-cell and those from bulk samples revealed that some CNA signatures observed in bulk tissue samples might not actually exist. Additionally, we have for the first time proposed the evolutionary trajectory of CNA signatures in HCC. These studies provide new insights into the mutation processes of HCC and important information for the in-depth study of CNA signatures and their potential clinical applications.

## Results

### Single-cell analysis revealed masked CNA profiles compared with bulk tissue CNA analysis

Each point mutation occurs on a single chromosome in a single cell, which gives rise to a lineage of cells bearing the same mutation. If that chromosomal locus is subsequently duplicated, any point mutation on this allele preceding the gain will subsequently be present on the two resulting allelic copies^25,26^. This process facilitates the accumulation of SBS without interference between mutations. Bulk tissue sequencing enables point mutation detection within a sample, reflecting the cumulative outcome of all subclones (Figure 1A). However CNA usually affect large proportion of genome DNA, when a CNA occurs within a single cell, the subsequent subclonal CNA further modify the original CNA profile, distorting its characteristic signature. Consequently, the CNA observed in bulk tissue is not an accurate reflection of the true CNA occurs in each subclones (Figure 1B). This phenomenon is particularly pronounced in tumors with high heterogeneity, such as HCC. To accurately explore the true state of CNA, we aggregated data from various HCC databases, covering different sequencing platforms and depths. Our collection comprised 1222 single-cell shallow whole-genome sequencing (sWGS) samples^24,27^, 160 bulk sWGS samples^28^, 178 bulk whole-genome sequencing (WGS) samples^29,30^, and 371 bulk SNP array samples^31^. All raw data were uniformly processed to derive the absolute copy number information of the samples (details provided in the Materials and Methods).

### Development of CNA signature analysis tool for both single-cell and bulk tissue samples

To identify single-cell CNA signatures, we developed a novel method for the extraction of CNA signatures. This method encompasses four principal aspects of CNA: absolute copy number, segment length, segment change, and segment shape (Figure 2). From these aspects, we further delineated 90 distinct features (Figure S1). These features were selected as hallmarks of previously reported genomic aberrations, including chromothripsis^32^, large-scale state transitions (LST)^33^, extrachromosomal circular DNA (ecDNA)^34^, and tandem duplications^35^. Following the computation of features for all samples, this feature matrix was processed using non-negative matrix factorization to identify CNA signatures (Figure 2). This method has previously been employed for extracting signatures of SBS^36,37^. The number of signatures extracted was determined using two parameters. First is the reconstruction error and the average Frobenius reconstruction error is reported. Second is the stability of signature extraction and the cosine similarity between the extracted signatures, and average silhouette width. In the single-cell samples, a total of seven CNA signatures were identified. In the analysis of bulk samples, seven CNA signatures were identified with sWGS and WGS data. Eight CNA signatures were determined using SNP array data (Figure S2).

**Figure 2.**
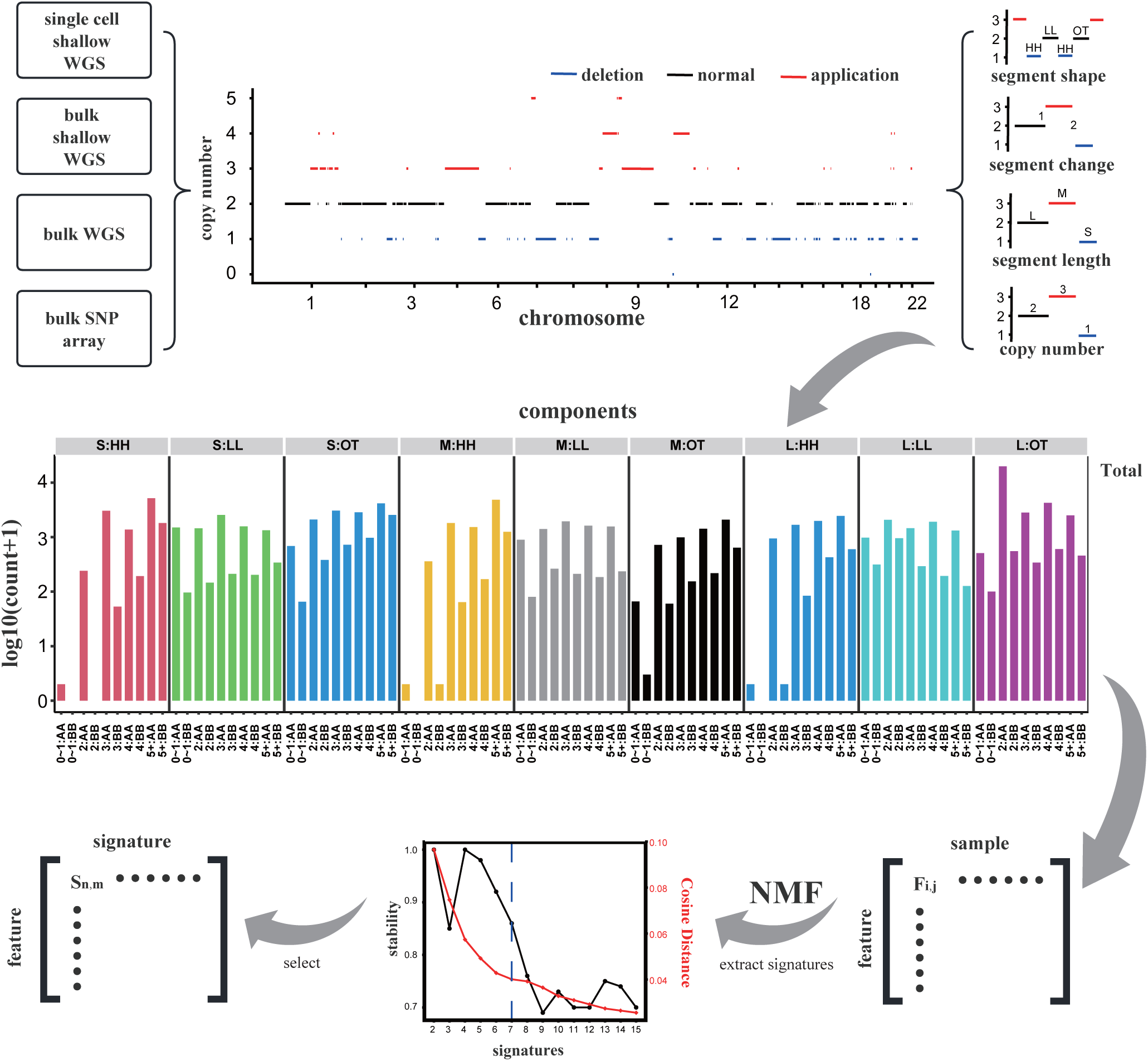
Workflow of CNA signature identification. The process of CNA signature extraction. A CNA feature matrix is generated from the absolute copy numbers obtained from different sequencing platforms and sequencing depths, and CNA signatures were extracted via NMF.

### Mutational processes for single-cell CNA signatures

Following determining CNA signatures, we initially assessed the similarity among different CNA signatures. The observed similarities between CNA signatures were low (Figure S3), suggesting that each CNA signature is relatively distinct (Figure 3A and Figure S4). Further, we compared the CNA signatures extracted from bulk tissue sWGS, WGS, and SNP array data, with those obtained from single-cell sWGS data. Our results indicate that the CNA signatures identified in single-cell shallow sequencing data can also be detected in other data types (Figure S5A-C), thus demonstrating the stability and reliability of our proposed CNA signature extraction method. However, specific CNA signatures, such as WGS_Sig5, WGS_Sig6, SNP_Sig3, and SNP_Sig4, did not have similar counterparts in single-cell CNA signatures. This phenomenon may be attributed to several factors. On the one hand, the limited number of patients represented in the single-cell samples may have resulted in the exclusion of certain sample-specific CNA signatures from the analysis. On the other hand, these CNA signatures may represent signals specific to bulk tissues rather than single cells.

**Figure 3.**
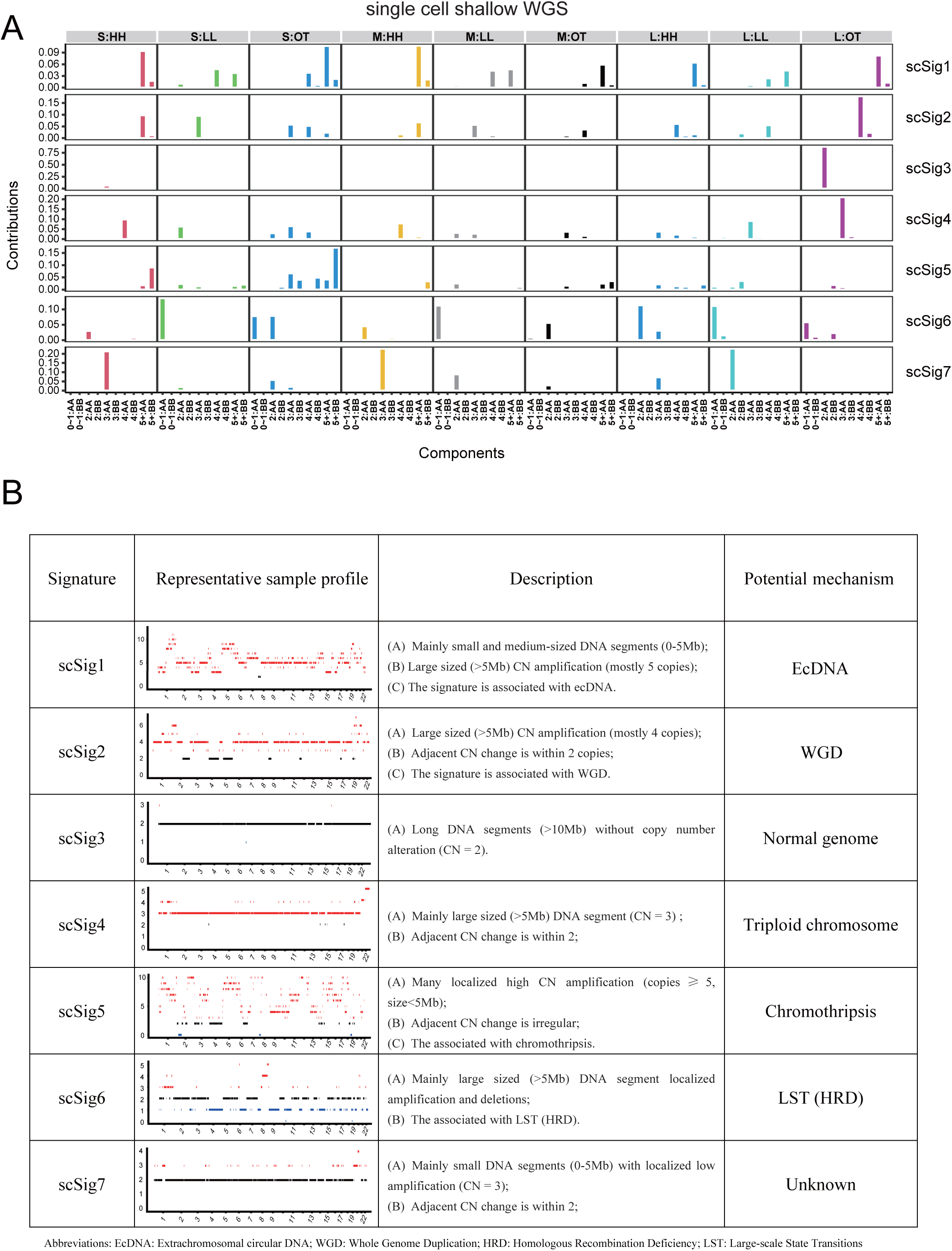
Profile and potential mutational processes of single-cell CNA signature (scSig). A, The CNA feature distribution of scSig, with the x-axis representing 90 types of CNA features and the y-axis indicating the contribution of each feature. B, Representative CNA profile, prominent features and potential mechanisms for each identified scSig.

Association analysis between CNA signatures and phenotypic data can reveal the underlying clinical and biological information for CNA signatures^8,11,15^. In the scSig1, we identified numerous genomic fragments smaller than 5Mb with high copy numbers, consistent with genomic variations induced by ecDNA^38–41^. Further associative analyses suggest these variations are attributable to ecDNA (Figure S6). Based on the analysis of representative samples of CNA signatures, we observed that the scSig3 pattern reflects characteristics of a normal genome, while the genome with the scSig4 pattern exhibits a clear triploid state, aligning with the CNS3 and CNS12 proposed by Tao et al^8^. Utilizing these analyses, we have summarized the potential etiologies of each CNA signature in single-cell data from HCC (Figure 3B).

### Single-cell CNA signature analysis reveals masked patterns in bulk tissue analysis

To further investigate the differences between single-cell CNA signatures and bulk tissue CNA signatures, following the methodology of Drews et al^7^, this study simulated CNA events, including early and late whole-genome duplication (early WGD, late WGD)^42,43^ and LST (associated with homologous recombination deficiency, HRD)^33^. The results indicate that single-cell CNA signatures can distinguish early WGD from late WGD, however bulk tissue CNA signatures fail to effectively differentiate early from late WGD events (Figure 4A). This finding reflects the advantag e of single-cell CNA signatures in finely distinguishing patterns of CNA. Following the genomic variation simulation method of Bartenhagen et al^44^, we simulated the formation of a series of CNA for comparative analysis, and the results also demonstrated that single-cell CNA signatures are more robust than bulk tissue CNA signatures in filtering out such simulated noise (Figure 4B and Figure S7).

**Figure 4.**
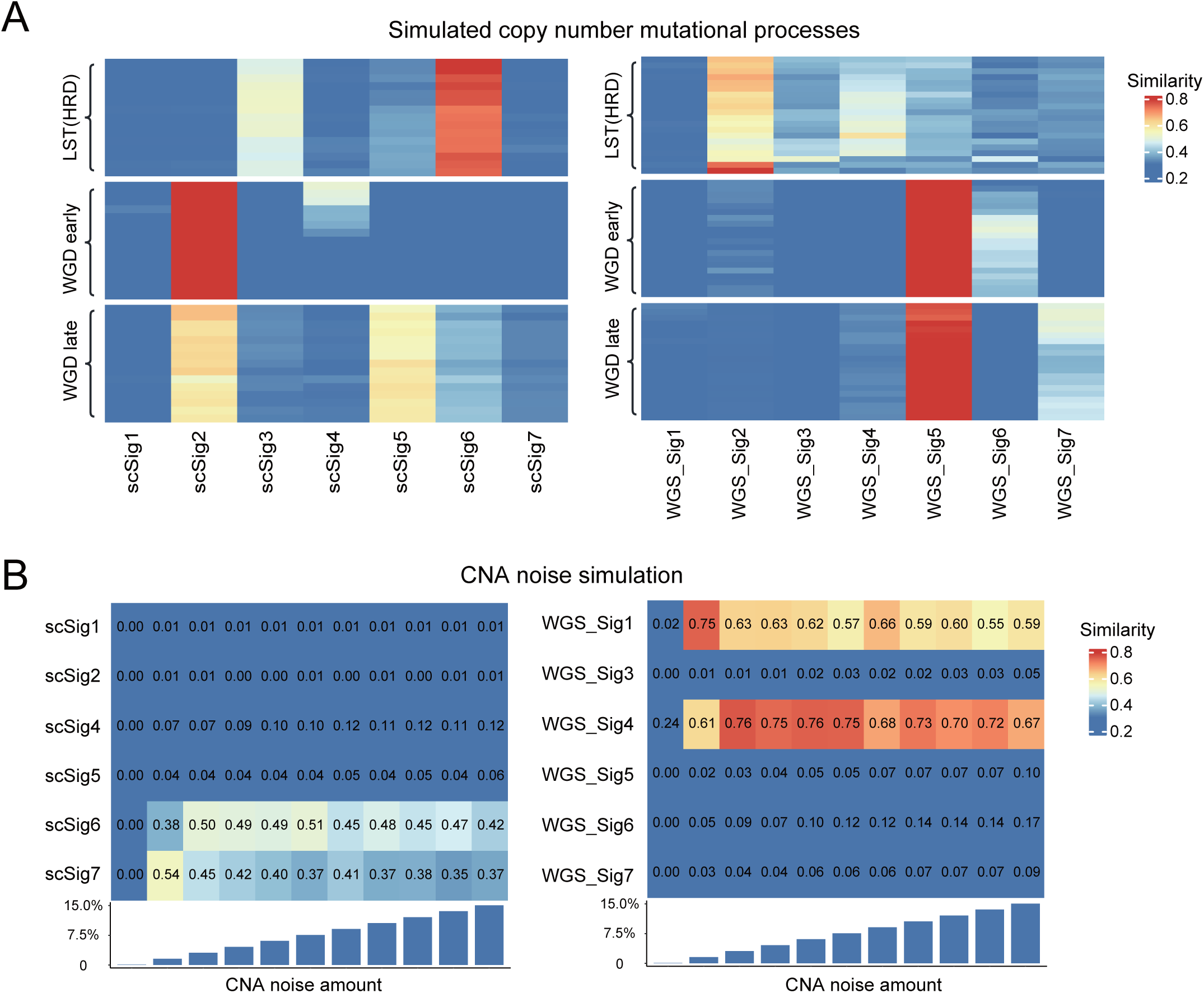
CNA mutational processes and noise simulation analysis suggest the robustness of scSig. A, Simulation of LST, early WGD, and late WGD events in twenty samples. The similarity between the CNA profile of each sample and the obtained CNA signature is calculated through cosine similarity. B, Simulate different levels of CNA noise (0∼15%), and calculate the cosine similarity between the CNA profile and the indicated CNA signature under each noise condition. Each noise level is simulated 100 times, and the mean similarity is used for plotting.

We employed the method of Collin Giguere et al^45^ to generate bulk tissue genomic sequencing data from corresponding single-cell data, and subsequently, absolute copy numbers were extracted (Figure S8). Then we fitted the generated bulk tissue CNA profiles with the single-cell CNA signatures. Comparing the differences reveals some inconsistencies between the CNA signatures of bulk tissue and corresponding single cells (Figures 5A-B). This inconsistency may be attributed to the CNA masking effects of bulk tissue samples. In the analysis of CNA patterns in individual samples, such as sample P04, we noted that the scSig4 constituted 96.7% in bulk tissue, whereas in corresponding single-cell samples, most cells predominantly exhibited scSig3 (Figures 5C-D and Figure S9). These findings suggest that in highly heterogeneous cancers like HCC, CNA signatures extracted from bulk tissue may not accurately reflect the true nature of the actual CNA in each cancer cells.

**Figure 5.**
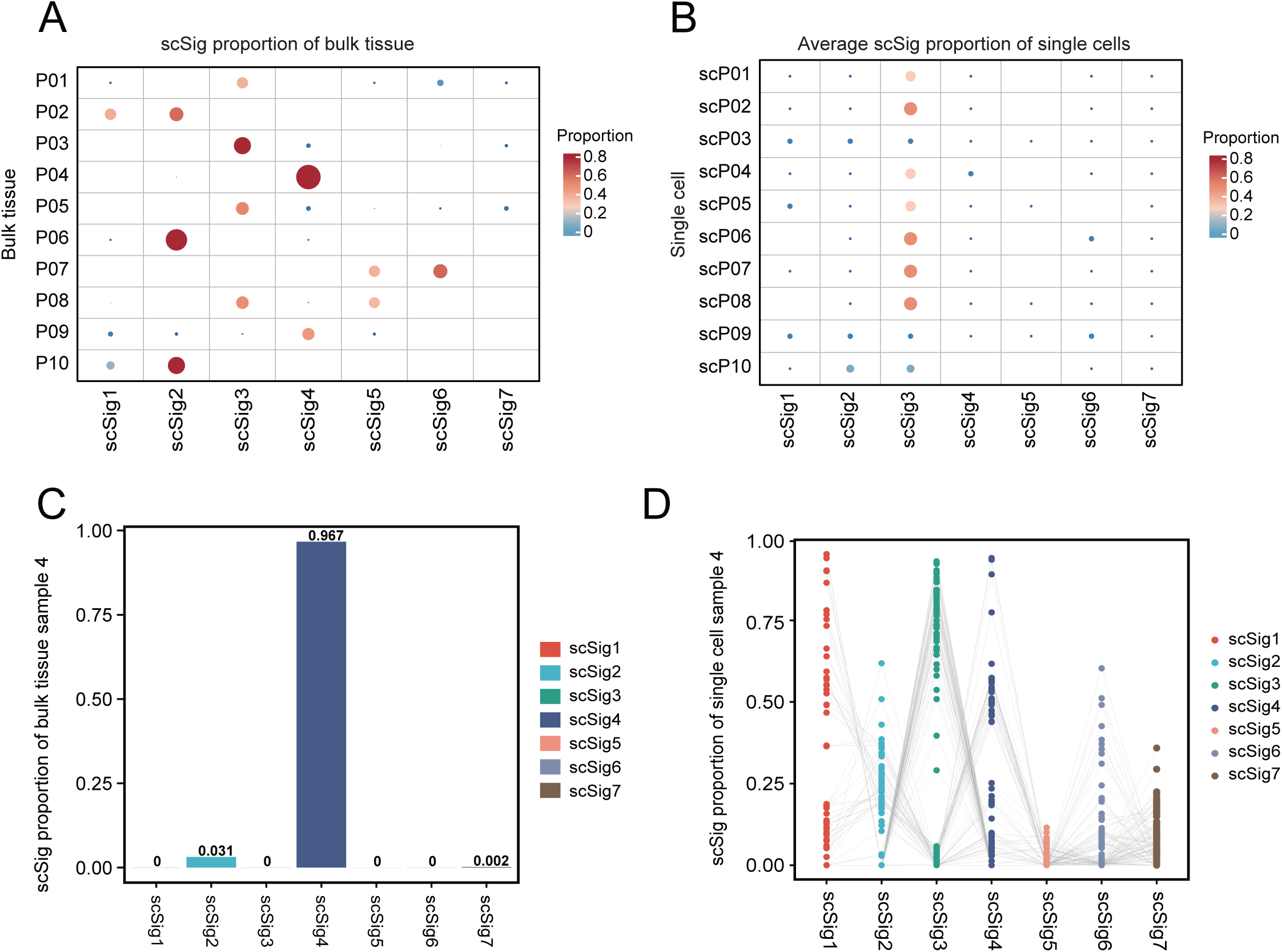
CNA signature comparison between single-cell and bulk tissue samples. A, Bulk tissue sample generated from corresponding single cells, and the distribution of scSig as a percentage of the total is presented. B, The mean proportion of each scSig across all cells within each sample is calculated, showing the average distribution of scSig within each sample. C, At the bulk tissue level, the distribution of scSig in sample P04, and scSig4 accounts for 96.7%. D, At the single-cell level, the distribution of scSig in sample P04, and scSig3 accounts for a relatively large proportion. Various scSig is present.

### Single cell CNA signature analysis reveals the evolutionary trajectory of HCC CNA signatures

We constructed a phylogenetic tree of CNA signatures to analyze the dynamic changes in the proportion of scSig across different single cells. The analysis revealed that scSig7 (characterized with small CNA) consistently appeared earlier in the evolutionary timeline. As the tumor progressed, the proportion of scSig1 and scSig2 (both are characterized with large CNA) gradually increased. These findings suggest that scSig7 happens in the early stages of tumorigenesis, while scSig1 and scSig2 are likely associated with the relatively late stages of HCC (Figure 6A and Figure S10). To further investigate the evolutional order of CNA signatures during HCC progression, we calculated the shared CNA breakpoints in individual cells to infer the potential evolutionary trajectories of tumor cells. The results indicated that scSig7 consistently appeared prior to other CNA signatures, while scSig1 and scSig2 were observed at later stages (Figure 6B and Figure S11). Additionally, we utilized the cancer CNA phylogenetic inference tool MEDICC2^46^ to construct a CNA evolutionary tree. Based on the pseudotemporal ordering of each cell in the evolutionary tree, we analyzed the dynamic changes of scSigs. Consistent with the previous findings, with scSig7 appearing at an earlier stage (Figure S12). These results elucidate the evolutionary trajectory of tumor genomic CNA signature, demonstrating that amplifications of small DNA fragments occur before large-scale genomic alterations in HCC evolution. The accumulation of these small fragment changes may affect the progression of the cell cycle, leading to chromosomal-level or genome-wide DNA alterations^47^.

**Figure 6.**
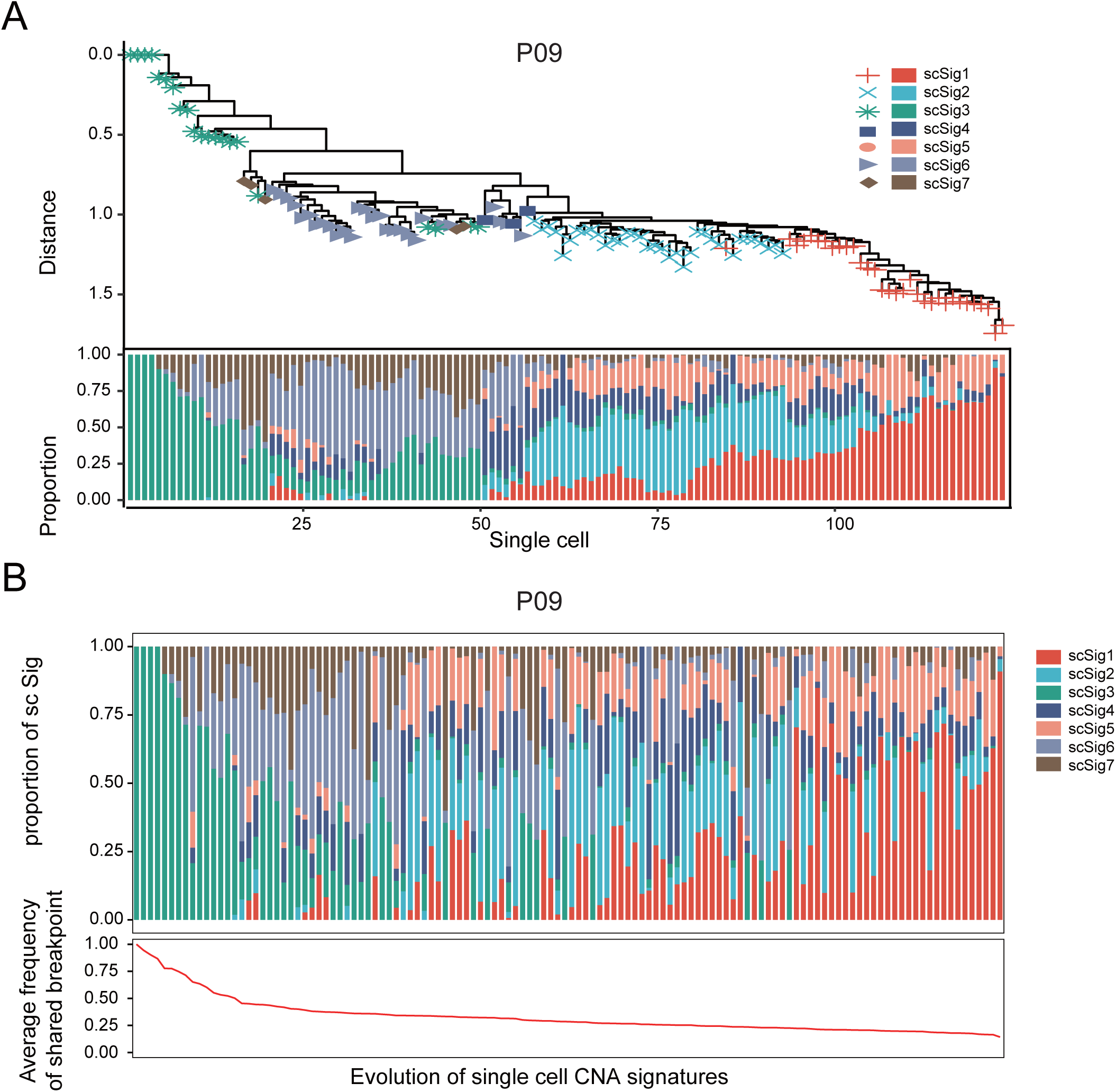
CNA signature evolution analysis in HCC. A, Evolutionary tree analysis of scSig. Calculate the distance between cells based on scSig, infer the evolutionary relationship based on the distance, and then draw an evolutionary tree. Symbols on each branch represent the most prevalent scSig within that lineage. B, Evolutionary trajectory analysis of scSig. The average number of shared breakpoints per cell within the sample is calculated, and cells are ranked according to the number of shared breakpoints. Cells with a higher ratio of shared breakpoints are positioned early in the clonal evolution, as subsequent subclones retain the breakpoints of earlier cells.

### Single-cell CNA signatures predict prognosis and drug response of HCC patients

Bulk tissue CNA signatures has been reported to be associated with cancer clinical outcomes^6,7,11,48^. To compare the prognostic impact of single-cell CNA signatures with those derived from WGS and SNP array CNA signatures, we analyzed HCC samples with survival data from the PCAWG and TCGA cohorts. We calculated the activity of scSig, WGS_Sig, and SNP_Sig in these samples through signature filtering. Across datasets from different technological platforms, scSig show more robust performance compared with WGS_Sig, and SNP_Sig as reflected in the consistent prognostic outcomes in different datasets (Figure 7A). In two independent datasets, scSig1 and scSig4 were associated with better prognosis, while scSig3 was related to poorer outcomes in the SNP dataset, with a similar trend observed in the WGS dataset (Figure 7B and Figure S13A-B). This finding suggests the superiority of single-cell CNA signatures over traditional bulk tissue CNA signatures in terms of HCC prognosis prediction.

**Figure 7.**
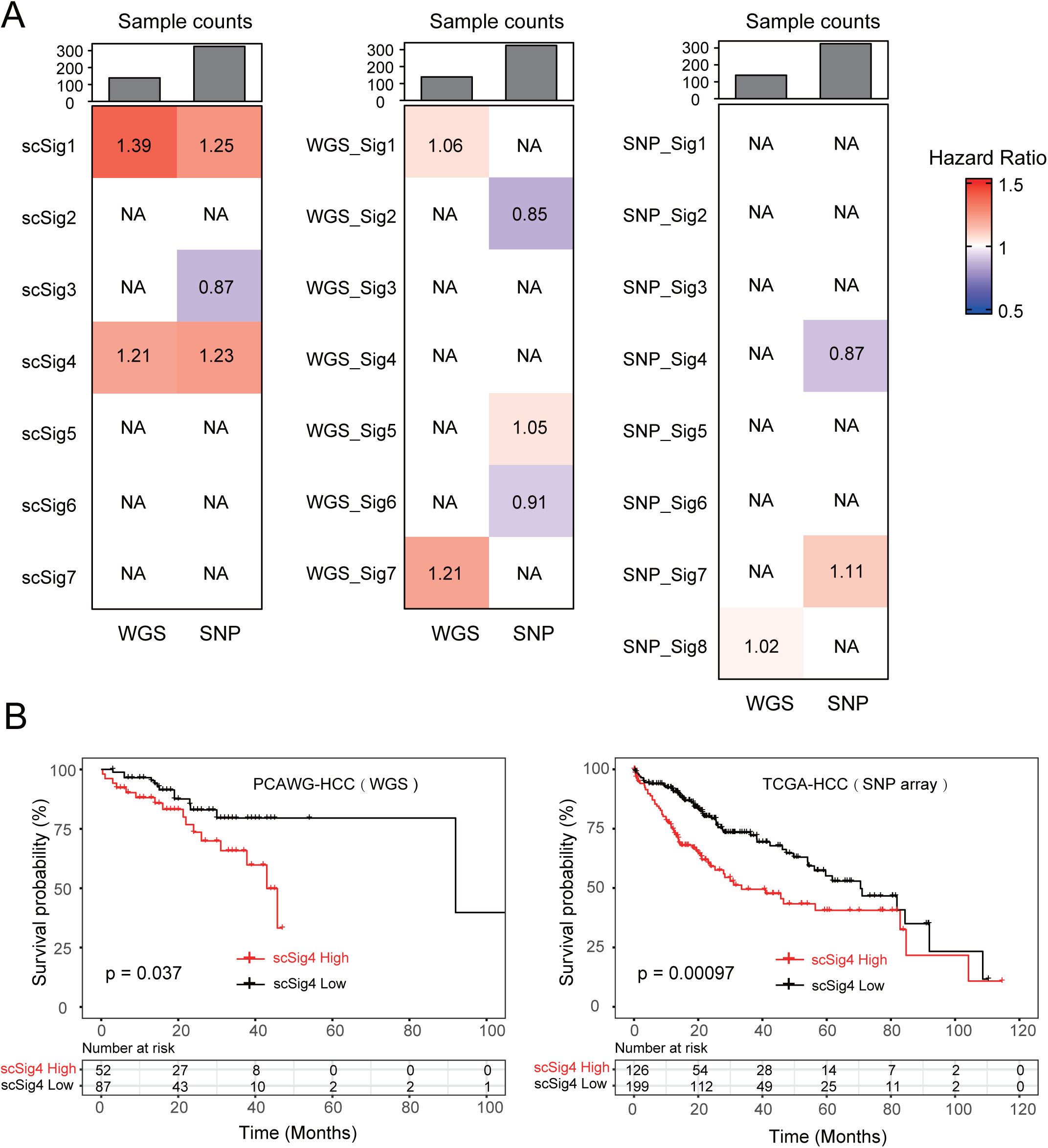
Single-cell CNA signatures show robust performance in prognosis prediction compared with CNA signatures derived from bulk tissues. A, The prognosis performance of scSig, WGS_Sig, and SNP_Sig in WGS and SNP array data was analyzed. Shown are the HR values of Signature that are significantly related to prognosis (FDR < 0.05), where HR > 1 indicates an association with poor prognosis, and HR < 1 indicates an association with better prognosis. B, The prognosis performance of the scSig4 in both WGS and SNP datasets. Consistent prognostic results in two different data sets.

We analyzed the correlations between drug sensitivity and CNA signatures. Our findings indicate that the sensitivity of HCC patients to the commonly used drug sorafenib is negatively correlated with the scSig7. Additionally, certain drugs demonstrated different responses to particular CNA signatures. Inhibitors of protein arginine methyltransferase (PRMT) have been found to impair tumorigenesis in HCC mouse models^49–51^. GSK591, a PRMT5 inhibitor, is regulated by MYC in HCC and has been identified as a direct target gene of MYC^52,53^. Patients carrying the scSig1 signature exhibited heightened sensitivity to GSK591, whereas they were non-responsive or showed resistance to other drugs (Figure S13C). Importantly, the scSig3 pattern, indicative of normal genomic features, was sensitive to most drugs, suggesting that early-stage tumors without significant CNAs have not yet developed substantial drug resistance. As genomic CNAs increase, tumors gradually acquire drug resistance. These results provide a new perspective for using single-cell CNA signatures in studying tumor drug resistance.

## Discussion

In this study, we have developed an analytical framework for the extraction of CNA signatures in single cells. This framework is adaptable across various experimental platforms and sequencing depths, effectively extracting CNA signatures.

CNA signatures have demonstrated their clinical value in a multitude of cancer studies^11,15^. Nevertheless, these existing CNA signature analyzes are all performed using bulk tissue samples, and single-cell CNA signature analysis is needed. We have systematically compared CNA signatures in single cells with those in bulk tissues of HCC. This study demonstrates that the majority of CNA signatures identified at the single-cell level correspond closely with those found in bulk tissue sequencing data of HCC. By employing existing methods to simulate CNA mutational processes and CNA noise, this research evaluated the reliability of different CNA signatures. The findings indicate that single-cell CNA signatures reveal potential CNA mutational processes with higher precision and exhibit greater stability in the presence of simulated noise. Despite the high heterogeneity of HCC, single-cell data accurately reflects the true state of CNA, as it is not affected by the mixing of subclones with different CNA.

This study is the first attempt to depict the dynamic changes of CNA signatures during HCC progression. We found that CNA signatures with small fragment amplifications, such as scSig7, typically appear in the early stages of HCC development. In contrast, CNA signatures indicative of chromosomal and genomic-level changes, such as scSig1 and scSig2, tend to emerge in the later stages. A reasonable explanation for this pattern is that amplification of small DNA fragments in the early stages disrupts normal cell cycle processes, subsequently leading to large-scale genomic variations, which contribute to the formation of complex large CNA patterns. However, these conclusions still require direct experimental evidence for further validation.

CNA is a complex biological process, and studies focusing on bulk tissues have already demonstrated its potential clinical utility. Nevertheless, the biological significance of certain CNA patterns remains to be elucidated, necessitating further in-depth investigation. In this study, we show that CNA signatures derived from single cells can effectively predict patient prognosis and drug sensitivity. Combining single-cell data with bulk tissue CNA data for integrated analysis could be a promising avenue for future research.

### Limitations of the study

From a single-cell perspective, we have identified some potential issues with bulk tissue CNA signatures. However, our study also has certain limitations. Although this study included data from 1,222 single cells, these data were derived from only 10 patients. The limited sample size might have led to the failure to detect all CNA features. Regarding the evolutionary trajectory analysis of CNA signatures, our analysis suggest that certain signatures consistently appear earlier in different samples. However, the complexity of the CNA process makes it challenging to establish an accurate evolutionary timeline and comprehensively describe the detailed evolutionary trajectory of CNA signatures. These analyses provide a direction for future research, and more experimental evidence is required for validation.

## Resource availability

### Lead contact

Further information and requests for resources should be directed to and will be fulfilled by the lead contact, Xue-Song Liu

### Materials availability

This study did not generate new materials.

### Data and code availability

Only publicly available data were used in this study, and data sources and handling of these data are described in the Materials and Methods. All codes required to reproduce the results reported in this manuscript are freely available at: https://github.com/XSLiuLab/single-cell-CNA-signature.

Further information is available from the corresponding author upon request.

## Method details

### Data collection and processing

We downloaded data from 1222 single-cell shallow whole-genome sequencing (0.4×) of HCC patient samples (totaling 10 patients) from the Genome Sequence Archive (GSA)^24,27^, under accession numbers HRA000094. Data from 160 HCC bulk samples with shallow whole-genome sequencing (0.5×) were also obtained from the International Cancer Genome Consortium (ICGC) database^28^. Moreover, absolute copy number data of whole-genome sequencing from 178 HCC bulk samples in the Pan-Cancer Analysis of Whole Genomes (PCAWG) cohort were acquired through the UCSC Xena platform^29,30^. Concurrently, SNP array data from 371 HCC bulk samples were sourced from The Cancer Genome Atlas (TCGA) database^31^.

For genome sequencing raw data (fastq) files, fastp was first used to perform quality control on the data, the BWA MEM algorithm was then applied using hg38 as reference genome^54^, and finally, the samtools tool was used to convert the SAM files into BAM files^55^. After obtaining the BAM files for all datasets, use the SortSam and MarkDuplicates commands in the Picard toolkit (https://broadinstitute.github.io/picard/) to sort BAM files and mark duplications for variant and copy number calling. Raw data processing the codes used can be found at Code availability.

### Absolute copy number calling

QDNAseq^56^ and ACE^57^ R packages were used to determine the copy number profiles from the low-coverage WGS BAM files. To determine the autosomal copy number profiles, we excluded both chromosome X&Y and mitochondrial DNA and the ploidy was adjusted using median bin segment value, which was the central assumption of ACE. For single-cell sequencing files, we used a ploidy penalty of 0.5 and lower-cellularity penalty of 1.0 to fit the “squaremodel()” function of ACE. For bulk tissue sequencing files, we used a ploidy penalty of 0.5 and lower-cellularity penalty of 0.5 to fit the “squaremodel()” function of ACE as per the author’s recommendation. ABSOLUTE^58^ was used to determine the copy number profiles from the SNP array files of TCGA. The code for copy number calling can be found in Code availability.

### CNA feature classification

Considering that shallow sequencing data cannot assess the LOH status of the genome, in order to describe the copy number variation of the liver cancer genome more accurately, we classify the copy number segments from four aspects, (1): copy number, which directly reflects the copy number amplification or deletion, divided into 6 states: 0, 1, 2, 3, 4, 5+, where 0 and 1 indicate that the genome fragment is missing, 2 indicates the normal genome state, and 3 and 4 indicate that the corresponding genome fragment is moderately amplified. , 5+ is considered highly amplified. (2): segment length, indicating the length of the copy number segment. The segment length of some small genomic variations such as ecDNA is mostly less than 5Mb^59,60^. Since most of the copy number variations produced by large scale transitions are greater than 10MB^61^, the copy number segment is length is divided into three types: S (5Mb<), M (5∼10Mb), and L (>10Mb). (3): segment change, indicating the change in copy number between two segments. Different mutations may imply that the mutations in adjacent segments come from different mechanisms. Divided into two types: AA (<=2) and BB (>2). (4): segment shape, the relationship between the copy number segment and two adjacent segments, divided into three types: HH (left High and right High), LL (left Low and right Low), and OT (other status).

### Deriving signatures

Copy number signatures were extracted from individual sample matrices using the gold standard tool SigProfiler v1.0.17 (https://github.com/AlexandrovLab/SigProfilerExtractor) with default parameters. Initially, the non-negative matrix factorization (NMF) algorithm was applied for the decomposition of the patient’s copy number feature matrix. Signature search intervals ranging from 3 to 12 were set, and NMF was run 200 times for each signature number with different random seeds to ensure result stability. For each NMF iteration, initialization was performed using random numbers, and iterations ranged from 10,000 to 1,000,000 until stable results were obtained. Two key parameters determined during this process were the stability of the signatures, as measured by silhouette analysis, and the average Frobenius reconstruction error. Under these criteria, seven signatures were identified as optimal at the single-cell level and were selected for further analysis.

### Copy number variation simulation

To simulate realistic chromosomal variation events, this study utilized the approach proposed by Drews et al^7^. We simulated WGD events by setting a tetraploid background prior to the insertion of other CNAs for early WGD, or by multiplying all copy number states by a factor following the placement of all CNAs for late WGD. Considering that most WGD events are early occurrences, the ratio of early to late WGD samples was set at 4:1^62^. The simulation of LST was based on their real breakpoint characteristics, employing a Poisson distribution for this simulation. Random copy number variations were simulated using the RSVSim^44^ tool, with hg38 as the reference genome. The simulations aimed to cover 0% to 15% of the genome. The length of each amplified and deleted segment was randomly set between 1kb and 5MB, distributed randomly across the entire genome, with all other parameters set to their default values. Calculating the degree of similarity between simulated samples and scSig using cosine similarity, a method widely used for assessing similarities among copy number profiles^8^.

### Phylogenic trees

We utilized the phangorn package in R to calculate the Maximum Parsimony Tree from the scSig matrix using the Parsimony Ratchet algorithm^63,64^. To ensure the accuracy of our analysis, samples that did not exhibit any CNAs were excluded. The branch lengths between cells were estimated using the ACCTRAN algorithm. The “ggtree” package in R was employed to visualize the phylogenetic trees^65^, where the predominant CNA signature were marked with distinct colors for easy visual differentiation.

### CNA signature evolution

To determine the temporal sequence of emergence for tumor cell subclones, we estimated the likely order by analyzing shared CNA breakpoints. Shared breakpoints refer to genomic CNAs that occur at the same positions across different subclonal cells within the same sample. Each subclone’s CNA may build upon the segments from prior CNA events, but these subclones are likely to retain the breakpoints from earlier CNAs.

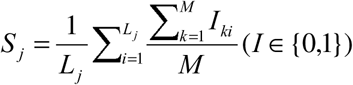

where S_j_ is the frequency of shared break point of cell j in the sample, M is the number of cells in the sample, L_j_ is the number of shared breakpoints in cell j, and I_ij_ represents the existence of the shared breakpoint I in cell j. I_ij_ is a binary variable. When the breakpoint i exists in the cell j, I_ij_ takes a value of 1; when the breakpoint i does not exist in the cell j, I_ij_ takes a value of 0.

### Survival analysis

In this study, we gathered survival data for samples from the TCGA (SNP array sequencing) and PCAWG (WGS sequencing) projects, encompassing 352 samples from TCGA and 139 samples from PCAWG. R package sigminer^15^ was utilized to fit each patient’s copy number feature matrix to scSig, SNP_Sig, and WGS_Sig, respectively. The association between CNA feature activity and overall survival was determined using univariate Cox proportional hazards models. For each Cox model, Hazard ratio were calculated and reported to indicate the direction and significance of the survival relationship. The Cox model is given by:

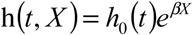

where t is the total survival time, h(t, X) is the hazard function given the presence of a prognostic variable X, and h0(t) represents the baseline hazard. Cox proportional hazards analysis was conducted using the R package ezcox^66^.

### Drug sensitivity analysis

Genomic information and drug sensitivity data for all HCC cell lines were downloaded from the GDSC database^67^. Following the previously introduced method, copy number variation data were obtained, and copy number feature matrices were calculated. Subsequently, using the R package sigminer, the copy number feature matrices of each cell line were fitted to scSig. Spearman correlation analysis was utilized to assess the relationship between the exposure values of scSig for each sample and IC50. The significance P-values obtained were corrected using the FDR method. Finally, the R package ComplexHeatmap^68^ was employed to draw correlation heatmap.

## Acknowledgments

We thank ShanghaiTech University High Performance Computing Public Service Platform for computing services. We thank multi-omics facility, molecular and cell biology core facility of ShanghaiTech University for technical help. This work is supported by cross disciplinary Research Fund of Shanghai Ninth People’s Hospital, Shanghai JiaoTong University School of Medicine (JYJC202227). Shanghai Science and Technology Commission (21ZR1442400), National Natural Science Foundation of China (82373149), and startup funding from ShanghaiTech University.

## Author Contributions

C.W. performed all the CNA signature analysis and drafted the manuscript under the supervision of X.-S. L. Z.T., W.C., N.W., J.W., J.S., T.W., D.Q., K.D., X.Z. T.L., L.Z., W.W., X.L., X.L., participated in cancer genome analysis and discussion. X.X. provided critical materials for this study and participated in critical project discussion and supervision. X.-S. L. conceived this study, supervised this study and wrote the manuscript with C.W.

## Supplement Figure Legends

**Figure S1. The distribution of the CNA features in different datasets.**

A, CNA feature matrix distribution of single-cell sWGS data. B, CNA feature matrix distribution of bulk sWGS data.

C, CNA feature matrix distribution of bulk WGS data.

D, CNA feature matrix distribution of bulk SNP array data.

**Figure S2. Determining the number of CNA signatures.**

The abscissa is the number of signatures, the y-axis on the left is the stability of each signature, and the right is the similarity between signatures. Signatures have stability and low similarity. A-D are single-cell sWGS data (A), bulk sWGS (B), bulk WGS (C), and bulk SNP array (D) data.

**Figure S3. Correlations between copy number signatures.**

Inter-correlation analysis of the profiles of different dataset CNA signatures. The numbers are cosine similarity values comparing each pair of CNA signatures. A-D are the signatures of scSig (A), sWGS_Sig (B), WGS_Sig (C), and SNP_Sig (D) datasets.

**Figure S4. Profiles of CNA signatures extracted from different types of datasets.**

The x-axis represents 90 types of CNA features and the y-axis indicates the contribution of each feature. A-C are the CNA feature matrices of bulk sWGS (A), bulk WGS (B), and bulk SNP array (C) data.

**Figure S5. Correlations of CNA signatures in different datasets**

A, Inter-correlations between the CNA signatures extracted in single-cell sWGS dataset and bulk WGS dataset. Cosine similarity values are reported for each comparison.

B, Inter-correlations between the CNA signatures extracted in single-cell sWGS dataset and bulk sWGS data dataset.

C, Inter-correlations between the CNA signatures extracted in single-cell sWGS dataset and bulk SNP array dataset.

D, Inter-correlations between the CNA signatures extracted in bulk WGS dataset and bulk SNP array dataset.

**Figure S6. Associations between ecDNA, chromothripsis and CNA signatures**

The Pearson correlation coefficient is shown, the circle’s size indicates the correlation level, and the color indicates significance. Significant associations with FDR adjusted P<0.05 are showed. scSig1 show the strongest correlation with ecDNA event, and scSig5 show the strongest correlation with chromothripsis event.

**Figure S7. CNA mutational processes and noise simulation analysis.**

A, Simulation of real CNA events, LST, early WGD, and late WGD were simulated in twenty samples. The similarity between the CNA profile of each sample and CNA signature is calculated through cosine similarity.

B, Simulate different levels of CNA noise (0∼15%), and calculate the cosine similarity between the CNA profile andCNA signature under each noise condition. Each noise level is simulated 100 times, and the mean similarity is used for plotting.

**Figure S8. CNA profiles of bulk tissue samples for each corresponding single-cell samples.**

Based on the raw data of single cells of each sample, the corresponding bulk tissue data is generated, and then the copy number is extracted. Black lines represent normal genome, blue lines represent deletions, and red lines represent amplifications.

**Figure S9. Comparison of scSig proportions between bulk samples and single-cell samples**

Distribution of scSig proportions in bulk tissue and single cells in samples P01, P05 and P09. There is a significant difference between scSig in bulk samples and scSig in single cells.

**Figure S10. The scSig evolutionary tree of single-cell samples.**

Evolutionary relationship of scSig. Calculate the distance between cells based on scSig, infer the evolutionary relationship based on the distance, and then draw an evolutionary tree. Symbols on each branch represent the most prevalent scSig within that lineage.

**Figure S11. scSig evolutionary trajectory analysis based on shared breakpoints.**

The average number of shared breakpoints per cell within the sample is calculated, and cells are ranked according to the ratio of shared breakpoints. Cells with a higher ratio of shared breakpoints are positioned early in the clonal evolution, as subsequent subclones retain the breakpoints of earlier cells.

**Figure S12. scSig evolutionary trajectory analysis based on the MEDICC2 algorithm.**

A, Single-cell CNA evolutionary trajectory of patient P09 calculated using the MEDICC2, with each branch representing a single-cell sample.

B, scSig evolution. Sorting of cells in the sample based on the pseudotemporal sequence calculated by MEDICC2, showing the proportion of each scSig in each single cell.

**Figure S13. Application of scSig signatures in HCC prognosis and drug sensitivity prediction.**

A, The prognosis performance of the scSig1 in both WGS and SNP datasets. Consistent prognostic results in two different datasets.

B, The prognosis performance of the scSig3 in both WGS and SNP datasets. Consistent prognostic results in two different datasets.

C, The relationship between scSig and drug sensitivity. Spearman correlation analysis was utilized to assess the relationship between the exposure values of scSig for each sample and IC50. The significance P-values obtained were corrected using the FDR method.

