## supplement figure for "Single-cell copy number alteration signature analysis reveals masked patterns and potential biomarkers for cancer"

Figure S1

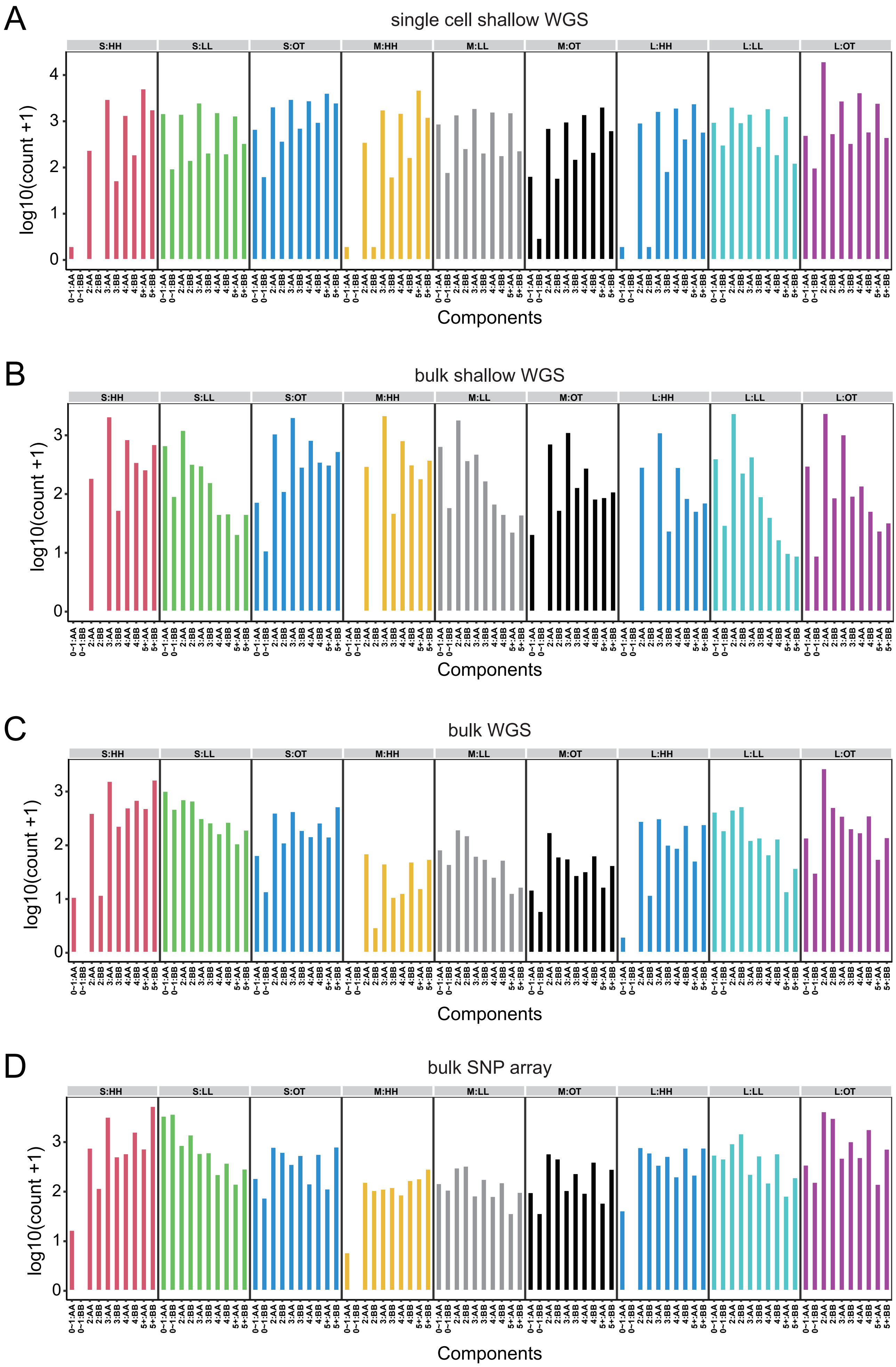
A
single cell shallow WGS
B
bulk shallow WGS
C
bulk WGS
D
bulk SNP array
S:HH
S:LL
S:OT
M:HH
M:LL
M:OT
L:HH
L:LL
L:OT
log10(count +1)
Components
0~1:AA
0~1:BB
2:AA
2:BB
3:AA
3:BB
4:AA
4:BB
5+:AA
5+:BB

**Figure S1. The distribution of the CNA features in different datasets.**

A, CNA feature matrix distribution of single-cell sWGS data.

B, CNA feature matrix distribution of bulk sWGS data.

C, CNA feature matrix distribution of bulk WGS data.

D, CNA feature matrix distribution of bulk SNP array data.

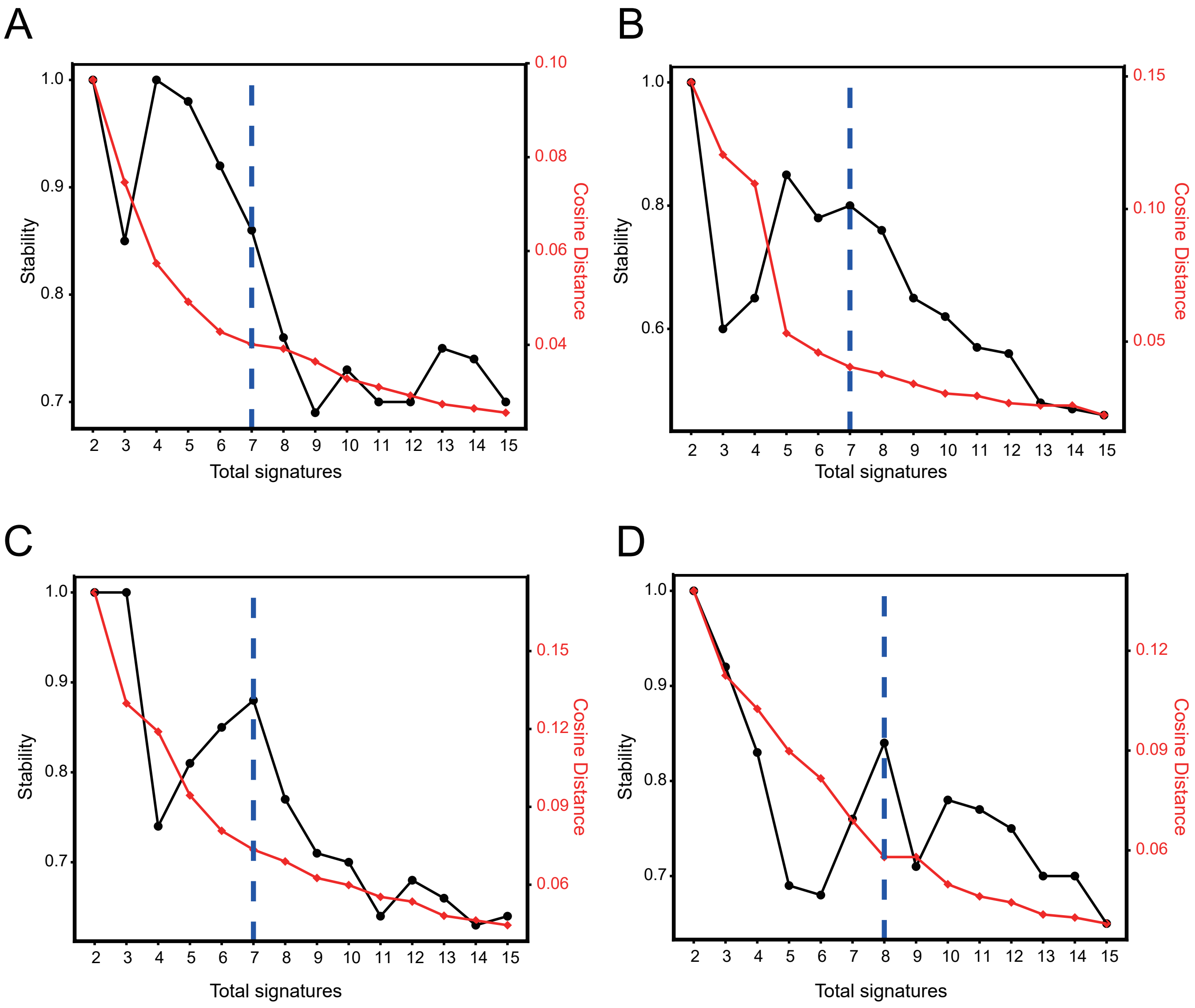


**Figure S2. Determining the number of CNA signatures.**

The abscissa is the number of signatures, the y-axis on the left is the stability of each signature, and the right is the similarity between signatures. Signatures have stability and low similarity. A-D are single-cell sWGS data (A), bulk sWGS (B), bulk WGS (C), and bulk SNP array (D) data.

Figure S3

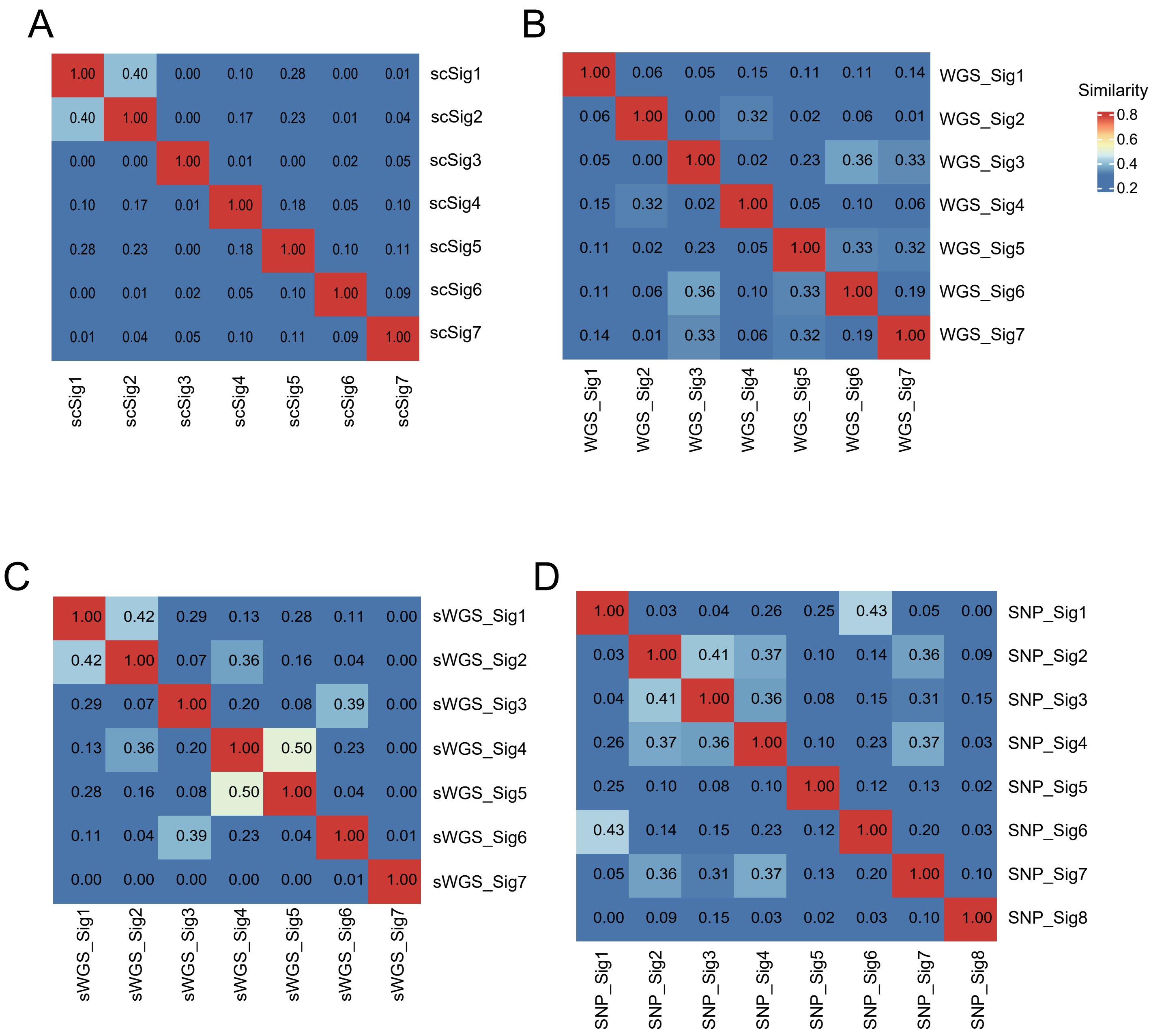


**Figure S3. Correlations between copy number signatures.**

Inter-correlation analysis of the profiles of different dataset CNA signatures. The numbers are cosine similarity values comparing each pair of CNA signatures. A-D are the signatures of scSig (A), sWGS_Sig (B), WGS_Sig (C), and SNP_Sig (D) datasets.

Figure S4

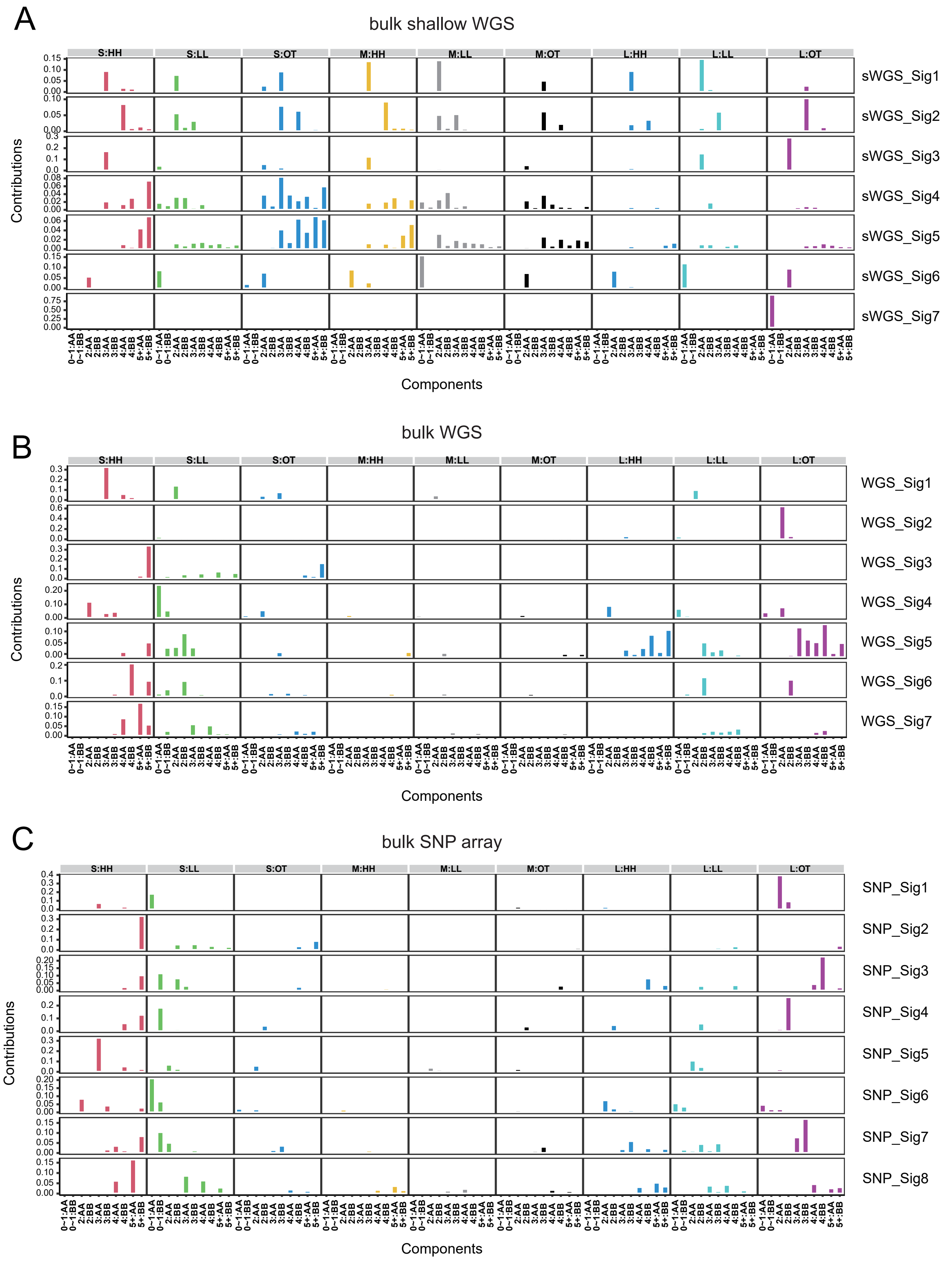
A
bulk shallow WGS
S:HH
S:LL
S:OT
M:HH
M:LL
M:OT
L:HH
L:LL
L:OT
sWGS_Sig1
sWGS_Sig2
sWGS_Sig3
sWGS_Sig4
sWGS_Sig5
sWGS_Sig6
sWGS_Sig7
Contributions
0−1:AA
0−1:BB
2:AA
2:BB
3:AA
3:BB
4:AA
4:BB
5+:AA
5+:BB
Components
B
bulk WGS
WGS_Sig1
WGS_Sig2
WGS_Sig3
WGS_Sig4
WGS_Sig5
WGS_Sig6
WGS_Sig7
C
bulk SNP array
SNP_Sig1
SNP_Sig2
SNP_Sig3
SNP_Sig4
SNP_Sig5
SNP_Sig6
SNP_Sig7
SNP_Sig8

**Figure S4. Profiles of CNA signatures extracted from different types of datasets.** The x-axis represents 90 types of CNA features and the y-axis indicates the contribution of each feature. A-C are the CNA feature matrices of bulk sWGS (A), bulk WGS (B), and bulk SNP array (C) data.

Figure S5

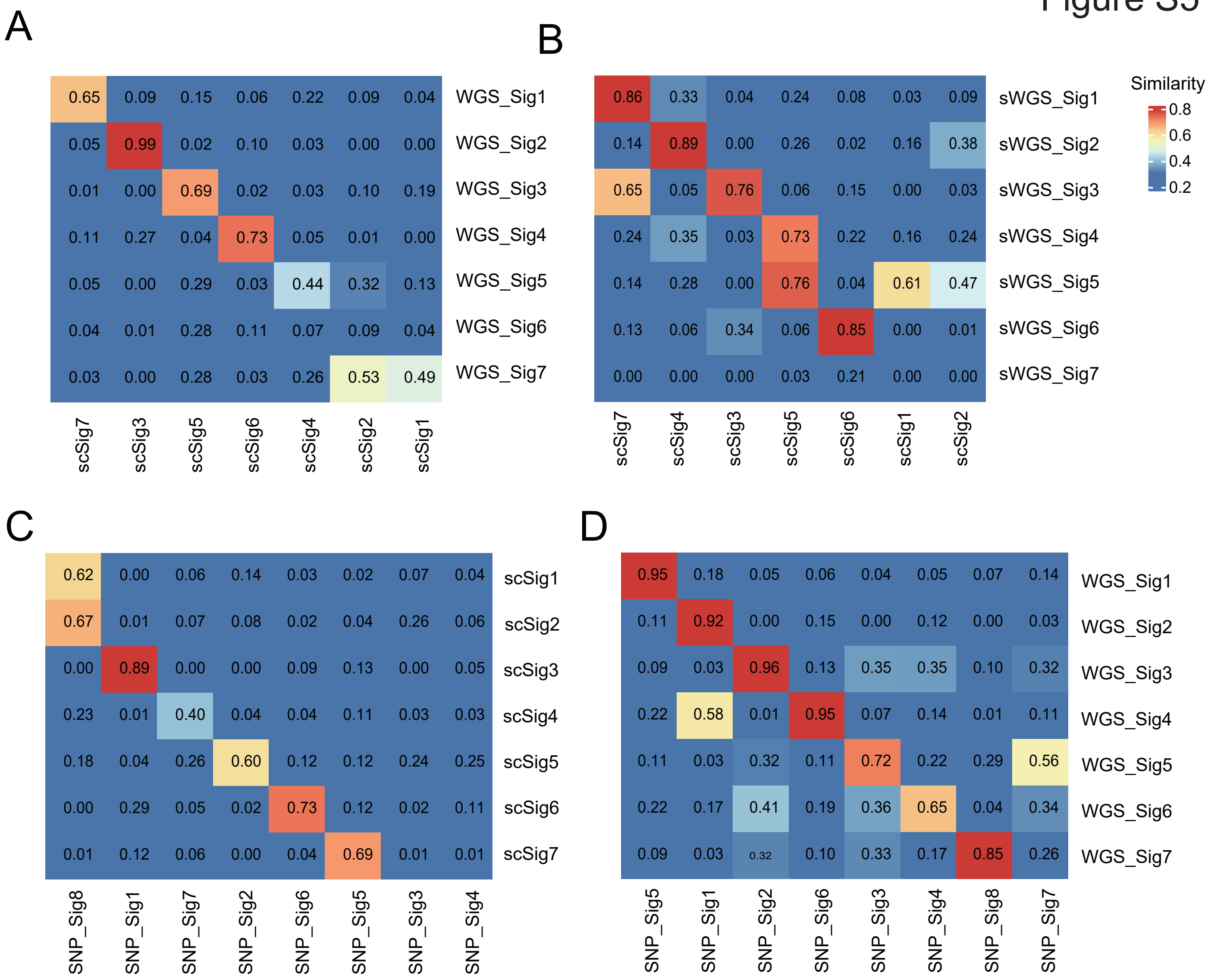


**Figure S5. Correlations of CNA signatures in different datasets**

A, Inter-correlations between the CNA signatures extracted in single-cell sWGS dataset and bulk WGS dataset. Cosine similarity values are reported for each comparison.

B, Inter-correlations between the CNA signatures extracted in single-cell sWGS dataset and bulk sWGS data dataset.

C, Inter-correlations between the CNA signatures extracted in single-cell sWGS dataset and bulk SNP array dataset.

D, Inter-correlations between the CNA signatures extracted in bulk WGS dataset and bulk SNP array dataset.

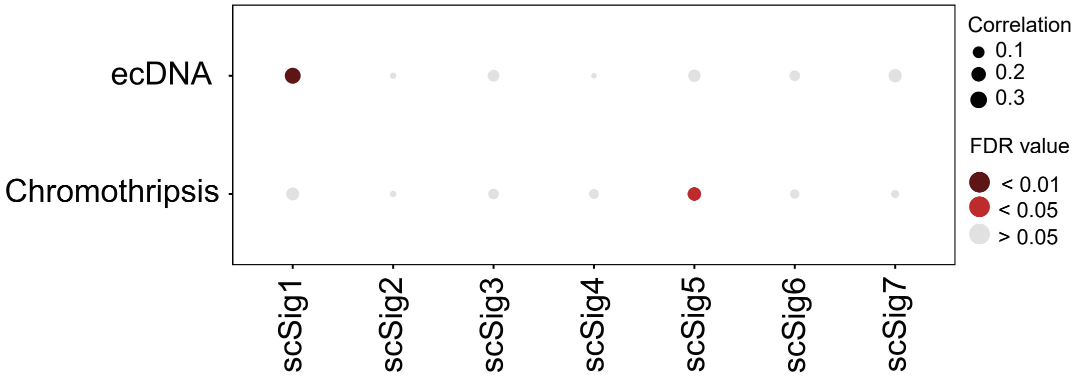


**Figure S6. Associations between ecDNA, chromothripsis and CNA signatures**

The Pearson correlation coefficient is shown, the circle's size indicates the correlation level, and the color indicates significance. Significant associations with FDR adjusted P<0.05 are showed. scSig1 show the strongest correlation with ecDNA event, and scSig5 show the strongest correlation with chromothripsis event.

Figure S7

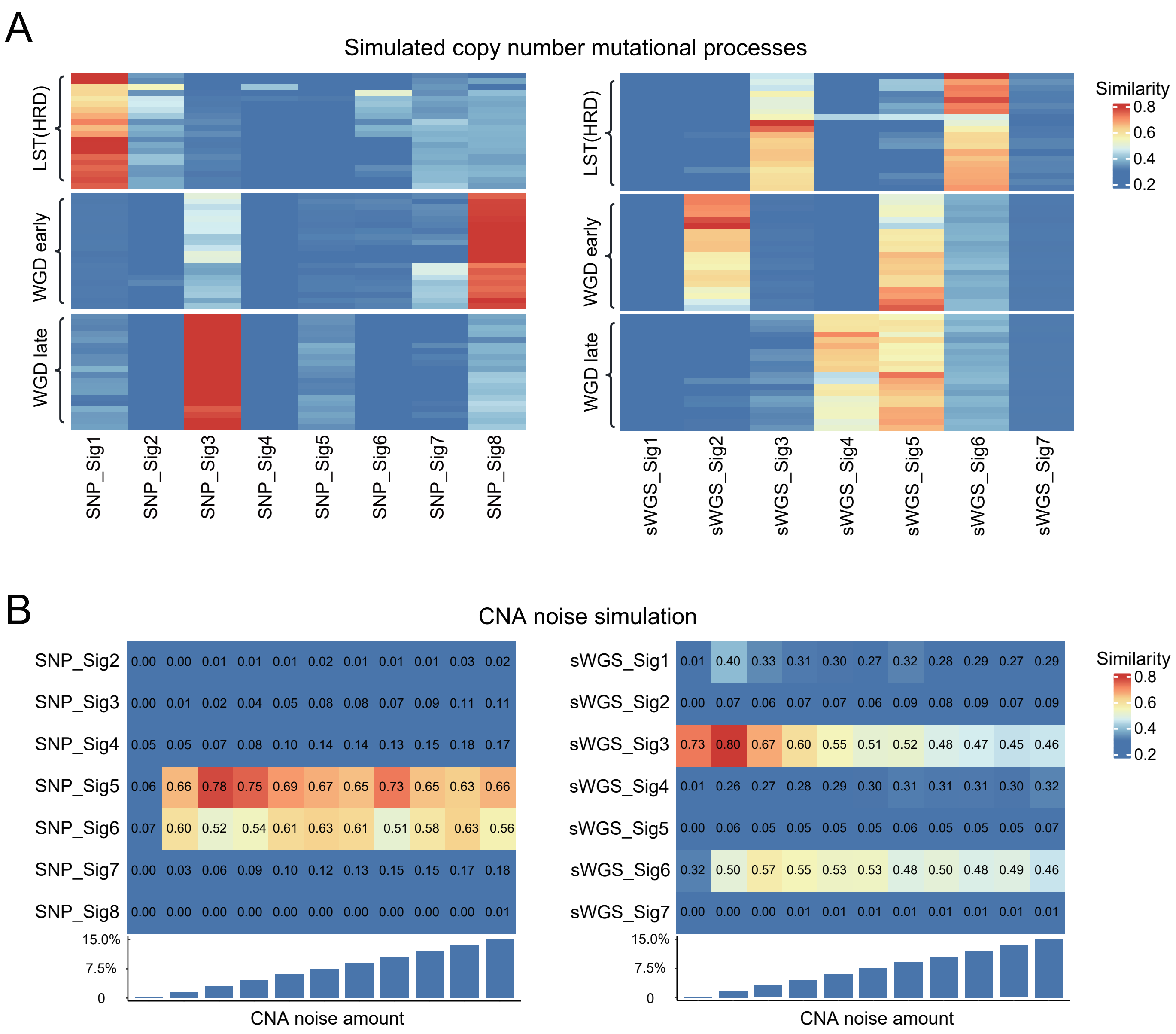


**Figure S7. CNA mutational processes and noise simulation analysis.**
A, Simulation of real CNA events, LST, early WGD, and late WGD were simulated in twenty samples. The similarity between the CNA profile of each sample and CNA signature is calculated through cosine similarity.
B, Simulate different levels of CNA noise (0~15%), and calculate the cosine similarity between the CNA profile andCNA signature under each noise condition. Each noise level is simulated 100 times, and the mean similarity is used for plotting.

Figure S8

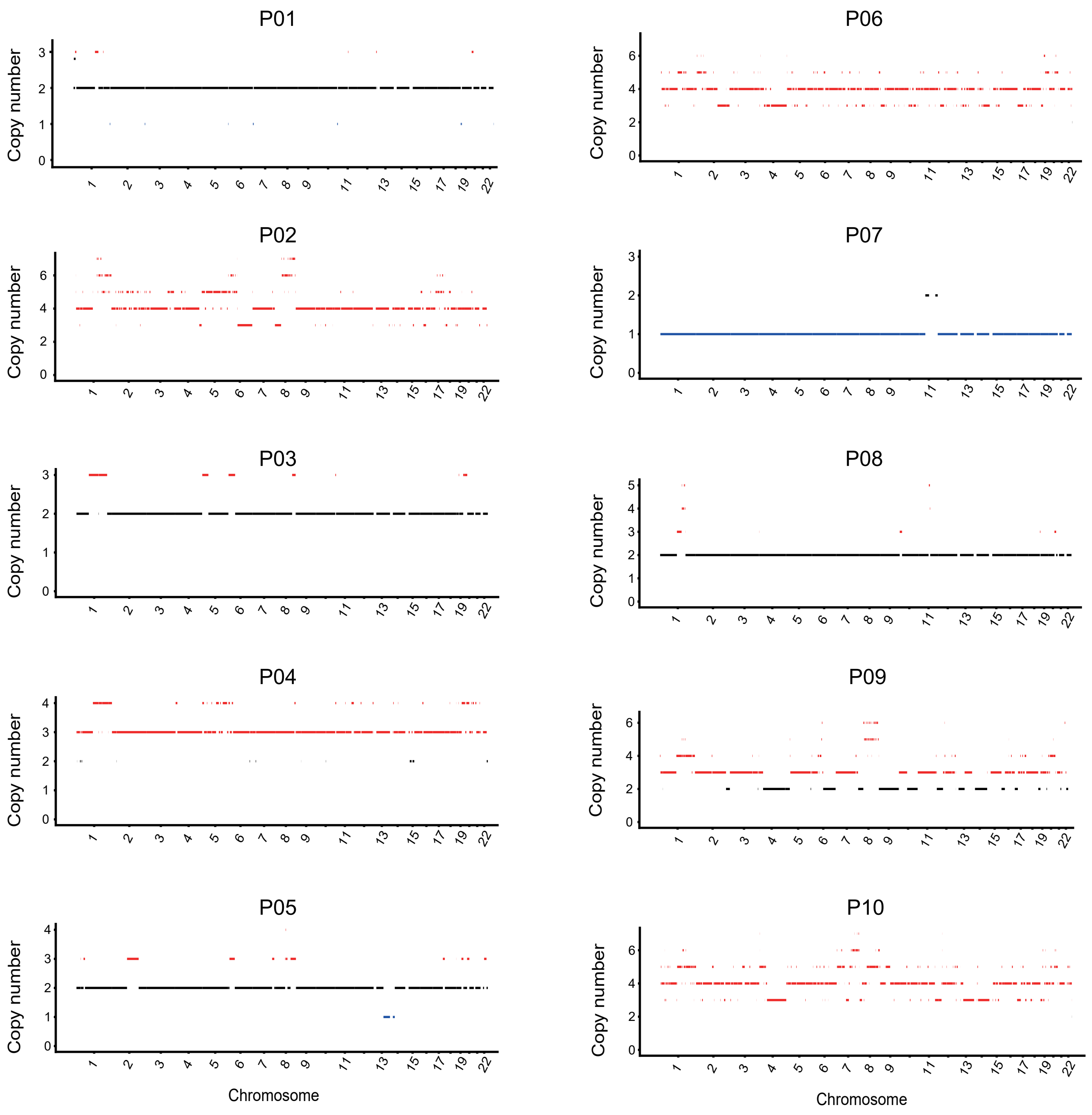


**Figure S8. CNA profiles of bulk tissue samples for each corresponding single-cell samples.**

Based on the raw data of single cells of each sample, the corresponding bulk tissue data is generated, and then the copy number is extracted. Black lines represent normal genome, blue lines represent deletions, and red lines represent amplifications.

Figure S9

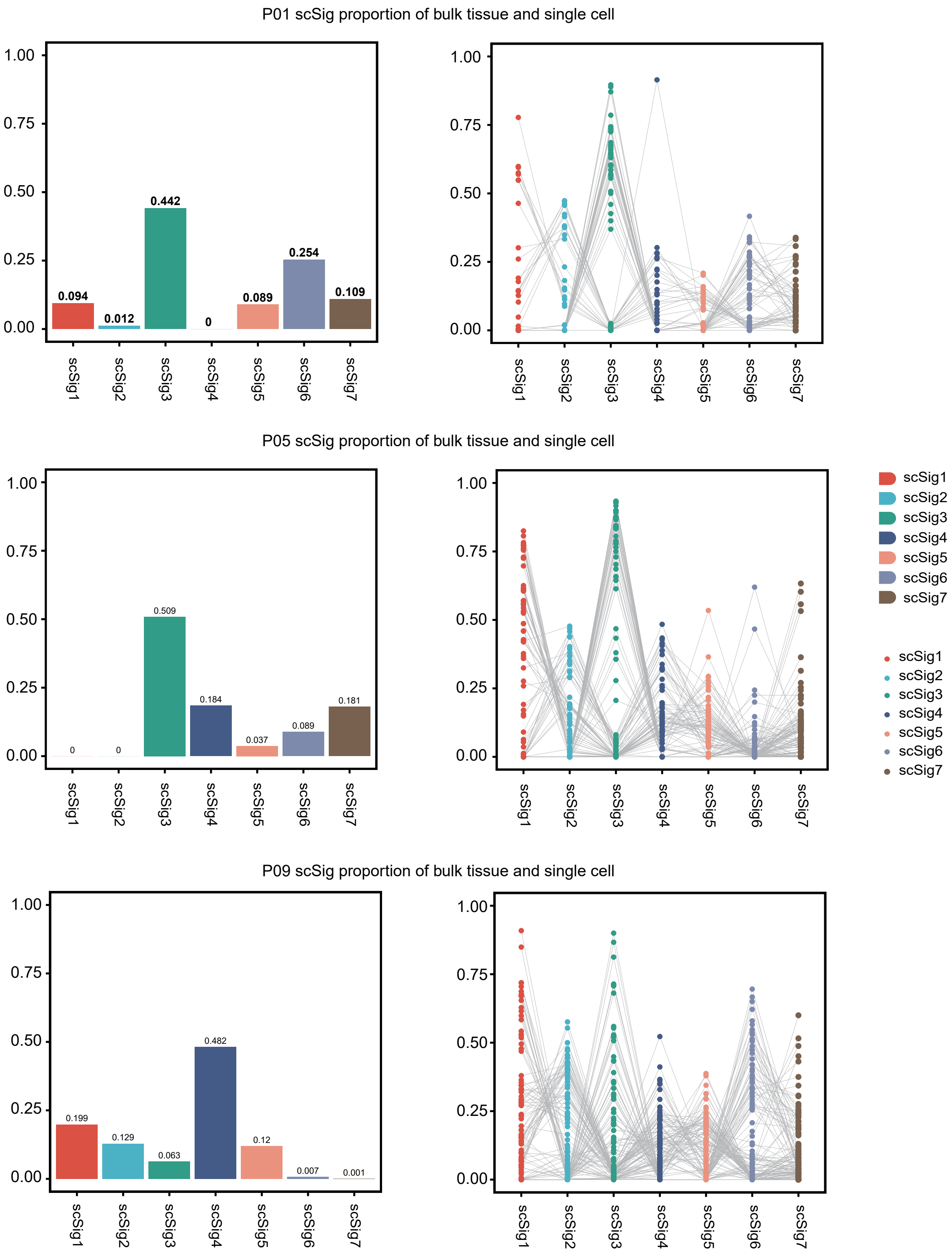


**Figure S9. Comparison of scSig proportions between bulk samples and single-cell samples**

Distribution of scSig proportions in bulk tissue and single cells in samples P01, P05 and P09. There is a significant difference between scSig in bulk samples and scSig in single cells.

### Figure S10

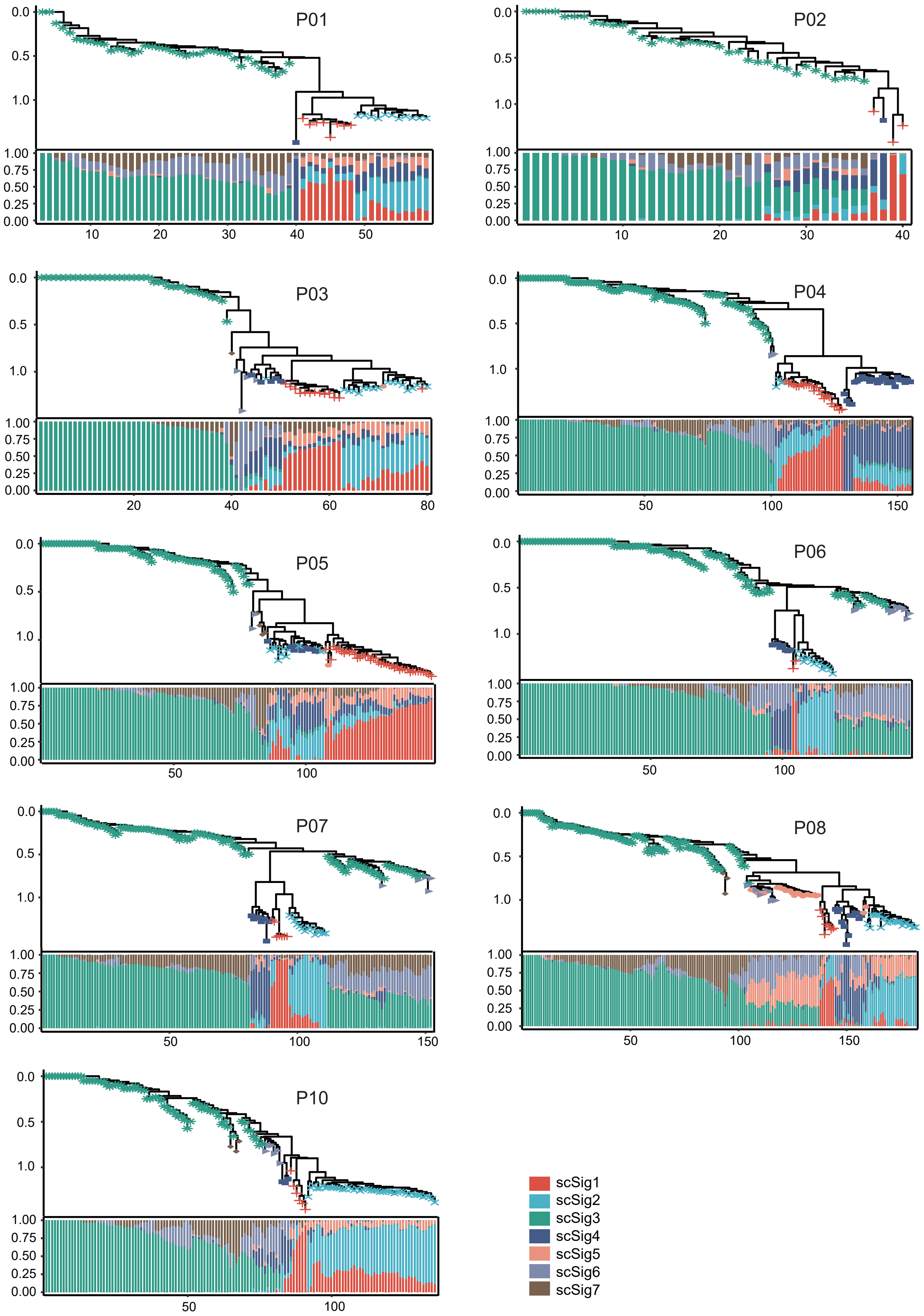
P01
P02
P03
P04
P05
P06
P07
P08
P10
0.0
0.5
1.0
1.00
0.75
0.50
0.25
0.00
scSig1
scSig2
scSig3
scSig4
scSig5
scSig6
scSig7

**Figure S10. The scSig evolutionary tree of single-cell samples.**

Evolutionary relationship of scSig. Calculate the distance between cells based on scSig, infer the evolutionary relationship based on the distance, and then draw an evolutionary tree. Symbols on each branch represent the most prevalent scSig within that lineage.

Figure S11

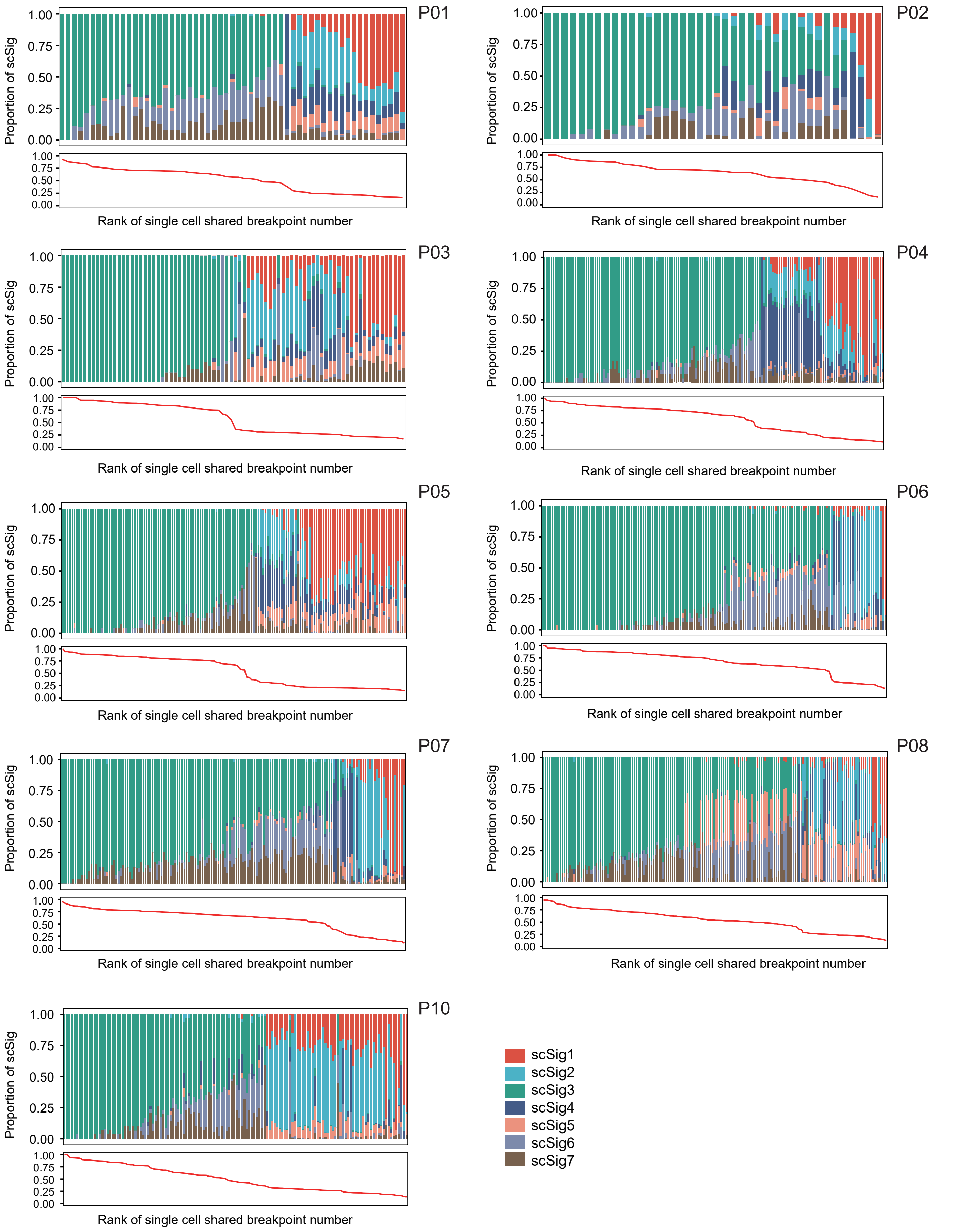
P01
P02
P03
P04
P05
P06
P07
P08
P10
Proportion of scSig
1.00
0.75
0.50
0.25
0.00
Rank of single cell shared breakpoint number
scSig1
scSig2
scSig3
scSig4
scSig5
scSig6
scSig7

**Figure S11. scSig evolutionary trajectory analysis based on shared breakpoints.**
The average number of shared breakpoints per cell within the sample is calculated, and cells are ranked according to the ratio of shared breakpoints. Cells with a higher ratio of shared breakpoints are positioned early in the clonal evolution, as subsequent subclones retain the breakpoints of earlier cells.

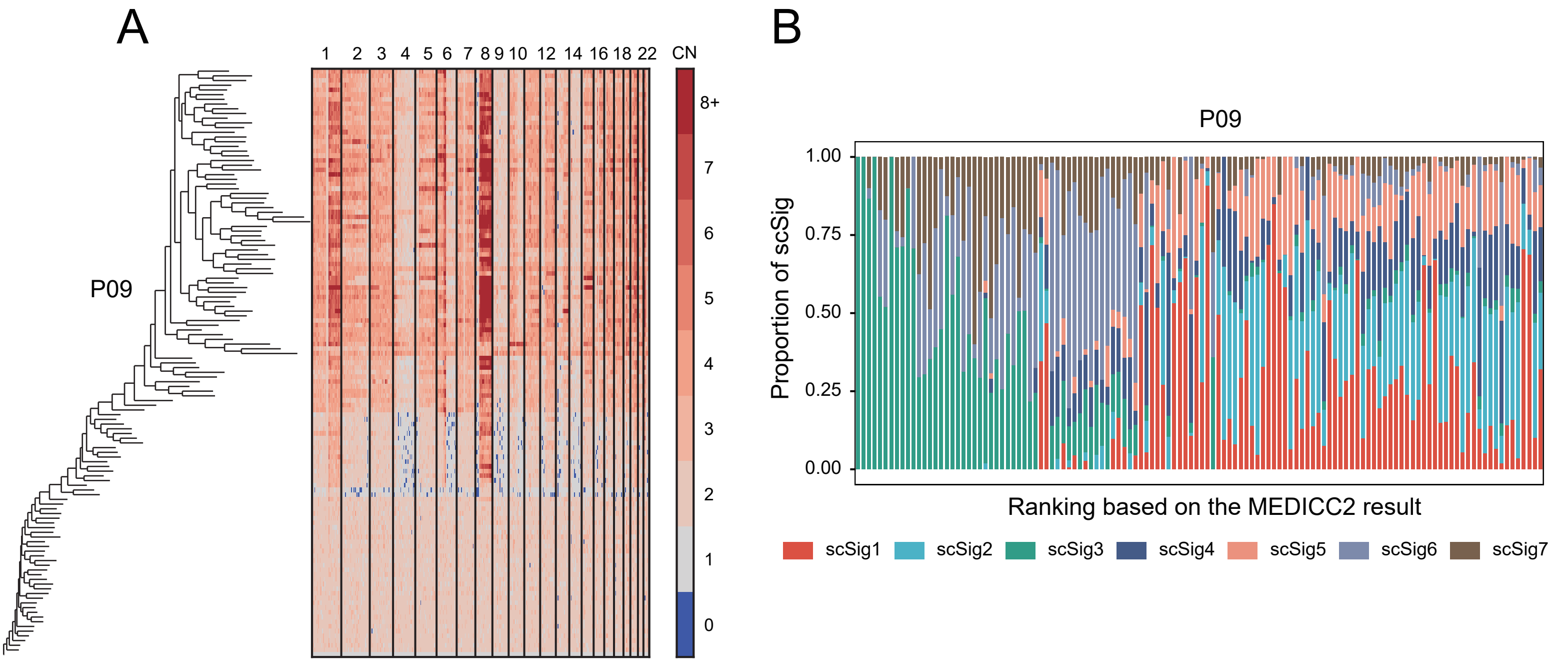


**Figure S12. scSig evolutionary trajectory analysis based on the MEDICC2 algorithm.**

A, Single-cell CNA evolutionary trajectory of patient P09 calculated using the MEDICC2, with each branch representing a single-cell sample.

B, scSig evolution. Sorting of cells in the sample based on the pseudotemporal sequence calculated by MEDICC2, showing the proportion of each scSig in each single cell.

Figure S13

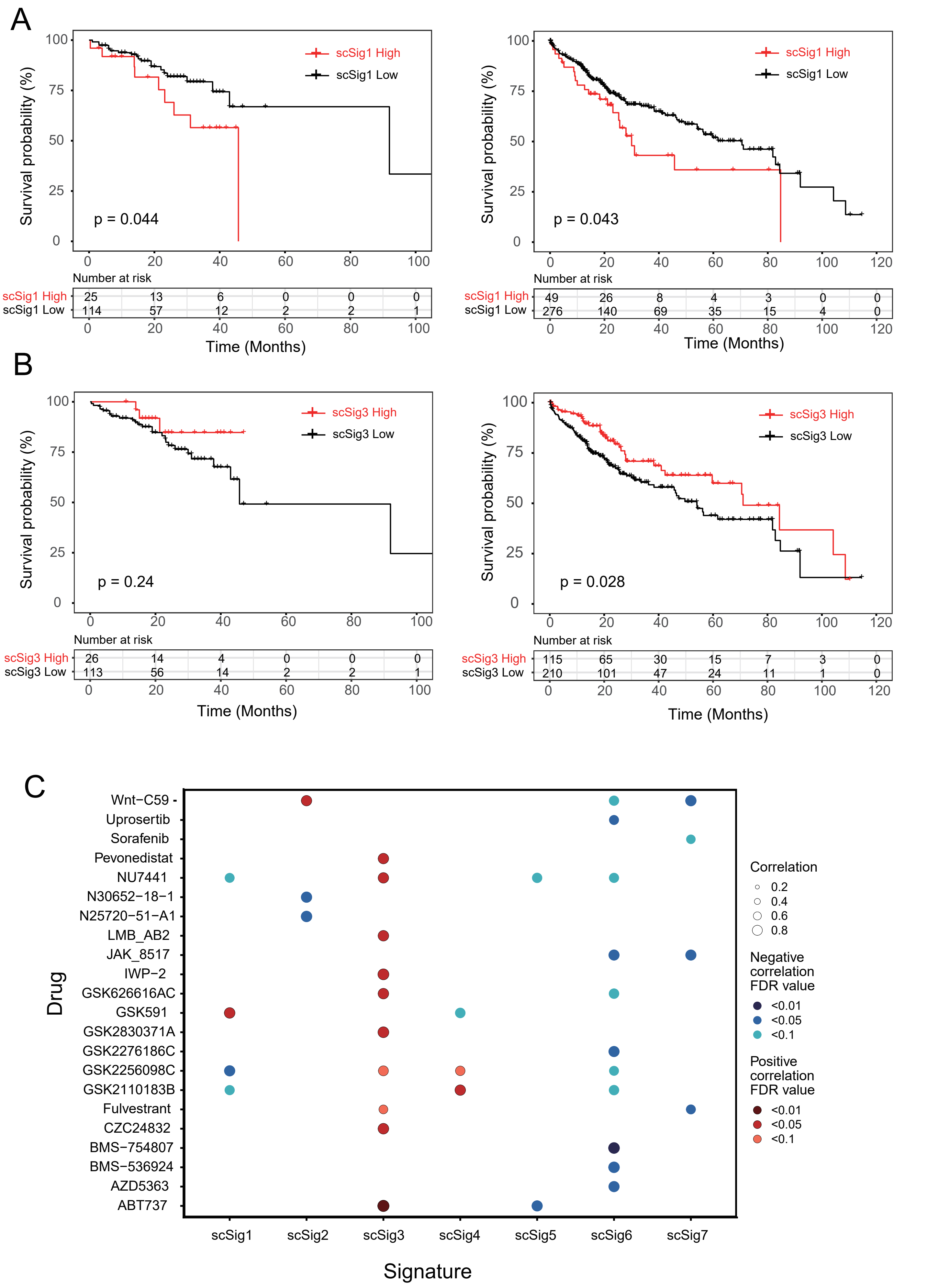

A
Survival probability (%)
scSig1 High
scSig1 Low
p = 0.044
p = 0.043
Number at risk
Time (Months)
B
scSig3 High
scSig3 Low
p = 0.24
p = 0.028
C
Drug
Wnt−C59
Uprosertib
Sorafenib
Pevonedistat
NU7441
N30652−18−1
N25720−51−A1
LMB_AB2
JAK_8517
IWP−2
GSK626616AC
GSK591
GSK2830371A
GSK2276186C
GSK2256098C
GSK2110183B
Fulvestrant
CZC24832
BMS−754807
BMS−536924
AZD5363
ABT737
scSig1
scSig2
scSig3
scSig4
scSig5
scSig6
scSig7
Signature
Correlation
0.2
0.4
0.6
0.8
Negative correlation FDR value
<0.01
<0.05
<0.1
Positive correlation FDR value
<0.01
<0.05
<0.1

**Figure S13. Application of scSig signatures in HCC prognosis and drug sensitivity prediction.**

A, The prognosis performance of the scSig1 in both WGS and SNP datasets. Consistent prognostic results in two different datasets.

B, The prognosis performance of the scSig3 in both WGS and SNP datasets. Consistent prognostic results in two different datasets.

C, The relationship between scSig and drug sensitivity. Spearman correlation analysis was utilized to assess the relationship between the exposure values of scSig for each sample and IC50. The significance P-values obtained were corrected using the FDR method.
